# Complementary cognitive roles for D2-MSNs and D1-MSNs during interval timing

**DOI:** 10.1101/2023.07.25.550569

**Authors:** R. Austin Bruce, Matthew A. Weber, Alexandra S. Bova, Rachael A. Volkman, Casey E. Jacobs, Kartik Sivakumar, Hannah R Stutt, Young-cho Kim, Rodica Curtu, Nandakumar S. Narayanan

## Abstract

The role of striatal pathways in cognitive processing is unclear. We studied dorsomedial striatal cognitive processing during interval timing, an elementary cognitive task that requires mice to estimate intervals of several seconds and involves working memory for temporal rules as well as attention to the passage of time. We harnessed optogenetic tagging to record from striatal D2- dopamine receptor-expressing medium spiny neurons (D2-MSNs) in the indirect pathway and from D1-dopamine receptor-expressing MSNs (D1-MSNs) in the direct pathway. We found that D2-MSNs and D1-MSNs exhibited distinct dynamics over temporal intervals as quantified by principal component analyses and trial-by-trial generalized linear models. MSN recordings helped construct and constrain a four-parameter drift-diffusion computational model in which MSN ensemble activity represented the accumulation of temporal evidence. This model predicted that disrupting either D2-MSNs or D1-MSNs would increase interval timing response times and alter MSN firing. In line with this prediction, we found that optogenetic inhibition or pharmacological disruption of either D2-MSNs or D1-MSNs increased interval timing response times. Pharmacologically disrupting D2-MSNs or D1-MSNs also changed MSN dynamics and degraded trial-by-trial temporal decoding. Together, our findings demonstrate that D2-MSNs and D1-MSNs had opposing dynamics yet played complementary cognitive roles, implying that striatal direct and indirect pathways work together to shape temporal control of action. These data provide novel insight into basal ganglia cognitive operations beyond movement and have implications for human striatal diseases and therapies targeting striatal pathways.

## Introduction

The basal ganglia have been heavily studied in motor control (Albin et al., 1989). These subcortical nuclei are composed of two pathways defined by medium spiny neurons (MSNs) in the striatum: the indirect pathway, defined by D2-dopamine receptor-expressing MSNs (D2- MSNs), and the direct pathway, defined by D1-dopamine receptor-expressing MSNs (D1- MSNs). It has been proposed that the indirect and direct pathways play opposing roles in movement (Alexander and Crutcher, 1990; Cruz et al., 2022; Kravitz et al., 2010), although recent work identified how these pathways can play more complex and complementary roles (Cui et al., 2013; Tecuapetla et al., 2016). MSNs receive dense input from both motor and cognitive cortical areas (Averbeck et al., 2014; Graybiel, 1997; Middleton and Strick, 2000), but the respective roles of D2-MSNs and D1-MSNs in cognitive processing are largely unknown.

Understanding basal ganglia cognitive processing is critical for diseases affecting the striatum such as Huntington’s disease, Parkinson’s disease, and schizophrenia (Andreasen, 1999; Hinton et al., 2007; Narayanan and Albin, 2022). Furthermore, pharmacological and brain stimulation therapies directly modulate D2-MSNs, D1-MSNs, and downstream basal ganglia structures such as the globus pallidus or subthalamic nucleus. Determining basal ganglia pathway dynamics during cognitive processes will help avoid side effects of current treatments and inspire novel therapeutic directions.

We studied striatal D2-MSNs and D1-MSNs during an elementary cognitive task, interval timing, which requires estimating an interval of several seconds. Interval timing provides an ideal platform to study cognition in the striatum because timing 1) requires cognitive resources including working memory for temporal rules and attention to the passage of time (Parker et al., 2013); 2) is reliably impaired in human striatal diseases such as Huntington’s disease, Parkinson’s disease, and schizophrenia (Merchant and de Lafuente, 2014); 3) requires nigrostriatal dopamine that modulates MSNs (Emmons et al., 2017; Gouvea et al., 2015; Matell et al., 2003; Mello et al., 2015; Monteiro et al., 2023; Wang et al., 2018); and 4) can be rigorously studied in animal models (Balci et al., 2008; Buhusi and Meck, 2005; Kim et al., 2017a; Parker et al., 2017, 2015). We and others have found that striatal MSNs encode time across multiple intervals by time-dependent ramping activity or monotonic changes in firing rate across a temporal interval (Emmons et al., 2017; Gouvea et al., 2015; Mello et al., 2015; Wang et al., 2018). However, the respective roles of D2-MSNs and D1-MSNs are unknown. Past work has shown that disrupting either D2-dopamine receptors (D2) or D1-dopamine receptors (D1) powerfully impairs interval timing by increasing estimates of elapsed time (Drew et al., 2007; Meck, 2006). Similar behavioral effects were found with systemic (Stutt et al., 2024) or focal infusion of D2 or D1 antagonists locally within the dorsomedial striatum (De Corte et al., 2019a). These data lead to the hypothesis that D2 MSNs and D1 MSNs have similar patterns of ramping activity across a temporal interval.

We tested this hypothesis with a combination of optogenetics, neuronal ensemble recording, computational modeling, and behavioral pharmacology. We use a well-described mouse-optimized interval timing task (Balci et al., 2008; Bruce et al., 2021; Larson et al., 2022; Stutt et al., 2024; Tosun et al., 2016; Weber et al., 2023). Strikingly, optogenetic tagging of D2- MSNs and D1-MSNs revealed distinct neuronal dynamics, with D2-MSNs tending to increase firing over an interval and D1-MSNs tending to decrease firing over the same interval, similar to opposing movement dynamics (Cruz et al., 2022; Kravitz et al., 2010; Tecuapetla et al., 2016). MSN dynamics helped construct and constrain a four-parameter drift-diffusion computational model of interval timing, which predicted that disrupting either D2-MSNs or D1-MSNs would increase interval timing response times. Accordingly, we found that optogenetic inhibition of either D2-MSNs or D1-MSNs increased interval timing response times. Furthermore, pharmacological blockade of either D2- or D1-receptors also increased response times and degraded trial-by-trial temporal decoding from MSN ensembles. Thus, D2-MSNs and D1-MSNs have opposing temporal dynamics yet disrupting either MSN type produced similar effects on behavior. These data demonstrate how striatal pathways play complementary roles in elementary cognitive operations and are highly relevant for understanding the pathophysiology of human diseases and therapies targeting the striatum.

## RESULTS

### Mouse-optimized interval timing

We investigated cognitive processing in the striatum using a well-described mouse-optimized interval timing task which requires mice to respond by switching between two nosepokes after a 6-second interval (Fig 1A; see Methods; (Balci et al., 2008; Bruce et al., 2021; Larson et al., 2022; Tosun et al., 2016; Weber et al., 2023)). In this task, mice initiate trials by responding at a back nosepoke, which triggers auditory and visual cues for the duration of the trial. On 50% of trials, mice were rewarded for nosepoking after 6 seconds at the designated ‘first’ front nosepoke; these trials were not analyzed. On the remaining 50% of trials, mice were rewarded for nosepoking at the ‘first’ nosepoke and then switching to the ‘second’ nosepoke; initial nosepokes at the second nosepoke after 18 seconds triggered reward when preceded by a first nosepoke. The first nosepokes occurred before switching responses and the second nosepokes occurred much later in the interval in anticipation of reward delivery at 18 seconds (Fig 1B-D). During the task, movement velocity peaked before 6 seconds as mice traveled to the front nosepoke (Fig 1E).

**Figure 1:**
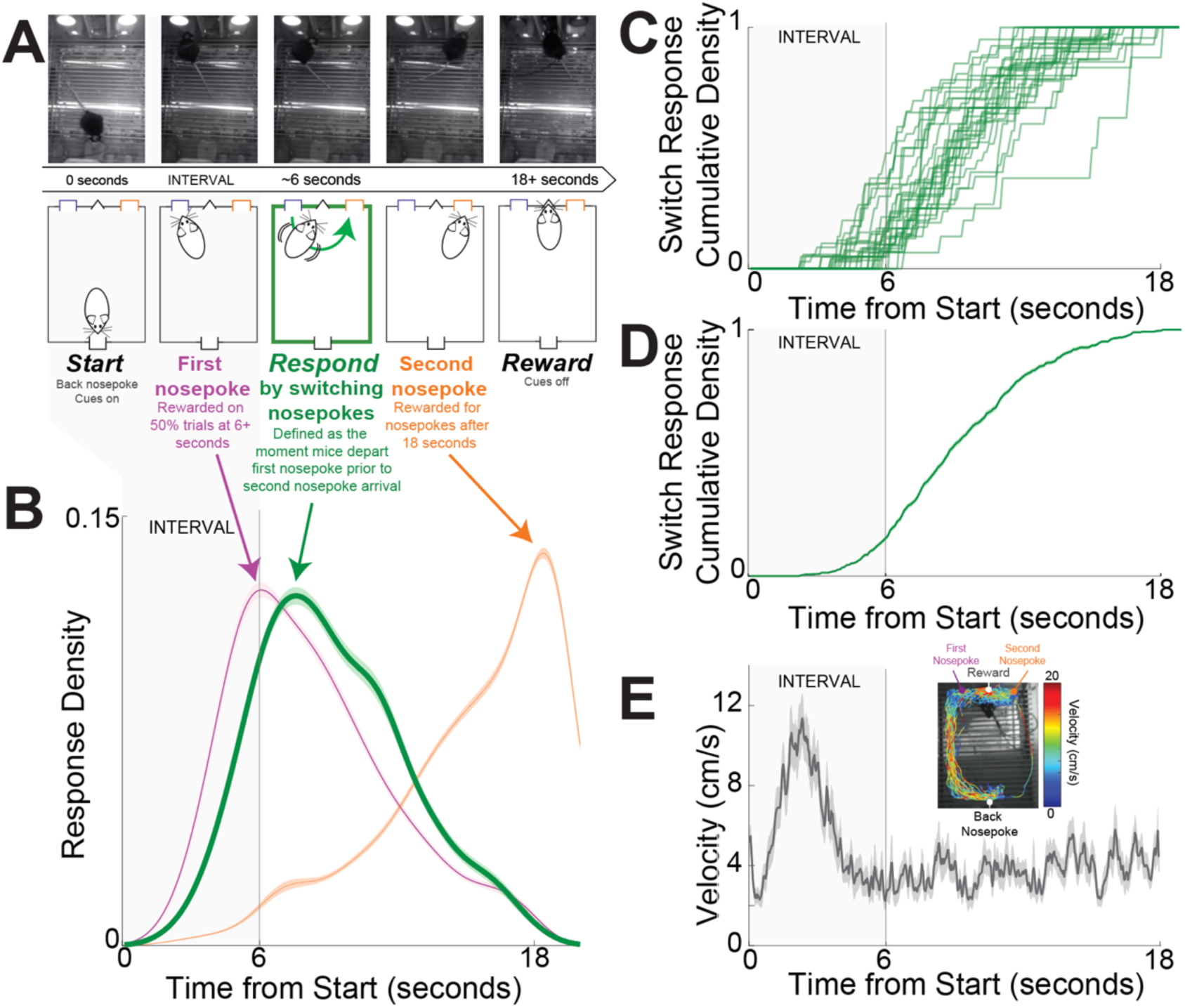
Mouse-optimized interval timing. A) We trained mice to perform an interval timing task in which they had to respond by switching nosepokes after a 6 second interval (in gray shade in all panels). Mice start trials by making a back nosepoke, which triggers an auditory and visual cue. On 50% of trials, mice were rewarded for a nosepoke after 6 seconds at the designated ‘first’ front nosepoke; these trials were not analyzed. On the remaining 50% of trials, mice were rewarded for switching to the ‘second’ nosepoke; initial nosepokes at the second nosepoke after 18 seconds triggered reward when preceded by a first nosepoke. *Switch response time* was defined as the moment mice depart the first nosepoke prior to second nosepoke arrival. Because cues are identical and on for the full trial on all trials, switch responses are a time-based decision guided explicitly by temporal control of action; indeed, mice switch nosepokes only if they don’t receive a reward at the first nosepokes after the 6 second interval. Top row - screen captures from the operant chambers during a trial with switch response. B) Response probability distribution from 30 mice for first nosepokes (purple), switch responses (green), and second nosepokes (orange). Responses at the first nosepoke peaked at 6 seconds, and switch responses peaked after 6 seconds. Because nosepoking at the second nosepoke was only rewarded after 18 seconds, second nosepokes tended to be highly skewed. Shaded area is standard error. C) Cumulative switch response density for each of 30 mice. D) Average cumulative switch response density; shaded area is standard error. E) DeepLabCut tracking of position during interval timing from a single mouse behavioral session revealed increased velocity after trial start and then constant velocity throughout the trial. Shaded area is standard error. In B-E, the 6 second interval is indicated in gray.

We focused on the *switch response time*, defined as the moment mice exited the first nosepoke before entering the second nosepoke. Switch responses are a time-based decision guided by temporal control of action because mice switch nosepokes only if nosepoking at the first nosepokes does not lead to a reward after 6 seconds (Fig 1B-E). Switch responses are guided by internal estimates of time because no external cue indicates when to switch from the first to the second nosepoke (Balci et al., 2008; Bruce et al., 2021; Tosun et al., 2016; Weber et al., 2023). We defined the first 6 seconds after trial start as the ‘interval’, because during this epoch mice are estimating whether 6 seconds have elapsed and if they need to switch responses.

In 30 mice, switch response times were 9.3 seconds (8.4 – 9.7; median (IQR)); see Table 1 for a summary of mice, experiments, trials, and sessions). We studied dorsomedial striatal D2-MSNs and D1-MSNs using a combination of optogenetics and neuronal ensemble recordings in 9 transgenic mice (4 D2-Cre mice switch response time 9.7 (7.0 – 10.3) seconds; 5 D1-Cre mice switch response time 8.2 (7.7 – 8.7) seconds; rank sum *p* = 0.73; Table 1).

**Table 1:**
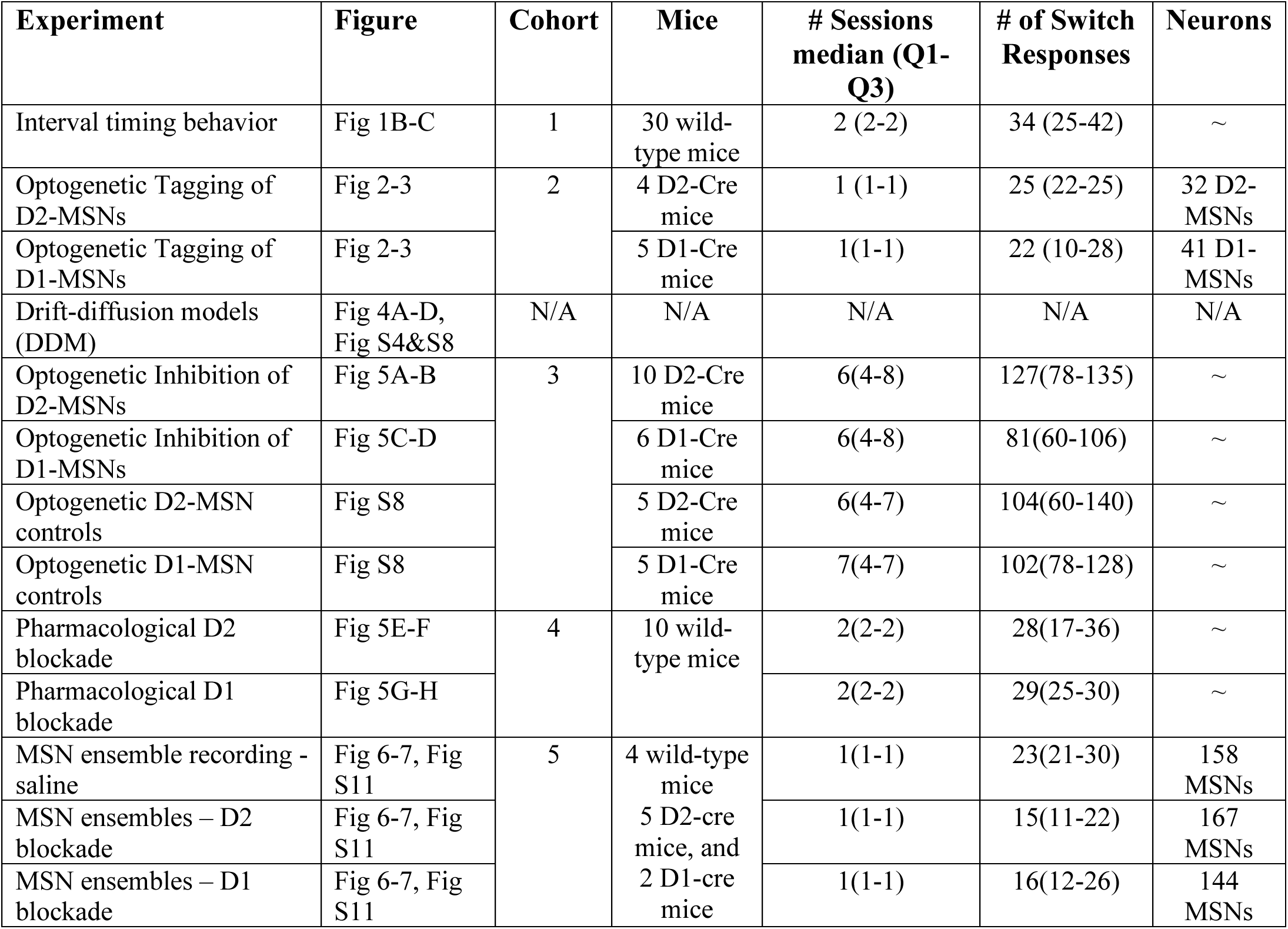
Summary of mice, sessions, # of switch trials, and MSNs.

### Opposing D2-MSN and D1-MSN dynamics

Striatal neuronal populations are largely composed of MSNs expressing D2-dopamine or D1-dopamine receptors. We optogenetically tagged D2-MSNs and D1-MSNs by implanting optrodes in the dorsomedial striatum and conditionally expressing channelrhodopsin (ChR2; Fig S1) in 4 D2-Cre (2 female) and 5 D1-Cre transgenic mice (2 female). This approach expressed ChR2 in D2-MSNs or D1-MSNs, respectively (Fig 2A-B; Kim et al., 2017a). We identified D2- MSNs or D1-MSNs by their response to brief pulses of 473 nm light; neurons that fired within 5 milliseconds were considered optically tagged putative D2-MSNs (Fig S1B-C). We tagged 32 putative D2-MSNs and 41 putative D1-MSNs in a single recording session during interval timing. There were no consistent differences in overall firing rate between D2-MSNs and D1- MSNs (D2-MSNs: 3.4 (1.4 – 7.2) Hz; D1-MSNs 5.2 (3.1 – 8.6) Hz; F = 2.7, *p =* 0.11 accounting for variance between mice). Peri-event rasters and histograms from a tagged putative D2-MSN (Fig 2C) and from a tagged putative D1-MSN (Fig 2D) demonstrate prominent modulations for the first 6 seconds of the interval after trial start. Z-scores of average peri-event time histograms (PETHs) from 0 to 6 seconds after trial start for each putative D2-MSN are shown in Fig 2E and for each putative D1-MSN in Fig 2F. These PETHs revealed that for the 6-second interval immediately after trial start, many putative D2-MSN neurons appeared to ramp up while many putative D1-MSNs appeared to ramp down. For 32 putative D2-MSNs average PETH activity increased over the 6-second interval immediately after trial start, whereas for 41 putative D1- MSNs, average PETH activity decreased. Accordingly, D2-MSNs and D1-MSNs had differences in activity early in the interval (0-5 seconds; F = 4.5, *p* = 0.04 accounting for variance between mice) but not late in the interval (5-6 seconds; F = 1.9, *p =* 0.17 accounting for variance between mice; Fig S2E). Examination of a longer interval of 10 seconds before to 18 seconds after trial start revealed the greatest separation in D2-MSN and D1-MSN dynamics during the 6-second interval after trial start (Fig S2). Strikingly, these data suggest that D2-MSNs and D1-MSNs might display distinct dynamics during interval timing.

**Figure 2:**
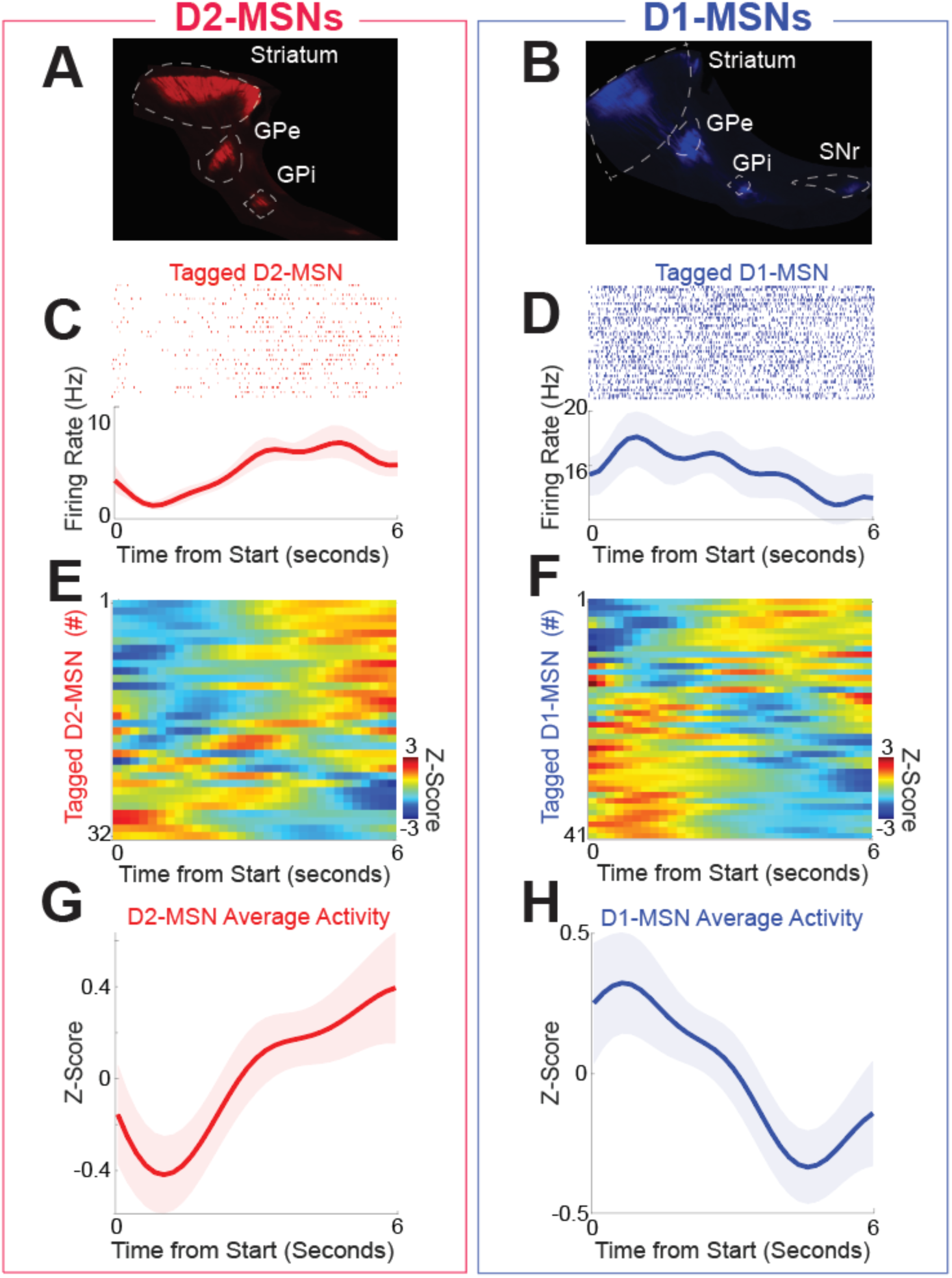
D2-MSNs and D1-MSNs have opposing dynamics during interval timing. A) D2-MSNs in the indirect pathway, which project from the striatum to the globus pallidus external segment (GPe; sagittal section) and internal segment (GPi) and B) D1-MSNs, which project from the striatum to the GPe, GPi, and substantia nigra (SNr; sagittal section). Peri-event raster C) from an optogenetically tagged putative D2-MSN (red) and D) from an optogenetically tagged putative D1-MSN (blue). Shaded area is the bootstrapped 95% confidence interval. E) Peri-event time histograms (PETHS) from all D2-MSNs and F) from all D1-MSNs. were binned at 0.2 seconds, smoothed using kernel-density estimates using a bandwidth of 1, and z-scored. Average activity from PETHs revealed that G) D2-MSNs (red) tended to ramp up, whereas H) D1-MSNs (blue) tended to ramp down. Shaded area is standard error. Data from 32 tagged D2-MSNs in 4 D2-Cre mice and 41 tagged D1-MSNs in 5 D1-Cre mice.

### Differences between D2-MSNs and D1-MSNs

To quantify differences between D2-MSNs vs D1-MSNs in Fig 2G-H, we turned to principal component analysis (PCA), a data-driven tool to capture the diversity of neuronal activity (Kim et al., 2017a). Work by our group and others has uniformly identified PC1 as a linear component among corticostriatal neuronal ensembles during interval timing (Bruce et al., 2021; Emmons et al., 2020, 2019, 2017; Kim et al., 2017a; Narayanan et al., 2013; Narayanan and Laubach, 2009; Parker et al., 2014; Wang et al., 2018). We analyzed PCA calculated from all D2-MSN and D1-MSN PETHs over the 6-second interval immediately after trial start. PCA identified time-dependent ramping activity as PC1 (Fig 3A), a key temporal signal that explained 54% of variance among tagged MSNs (Fig 3B; variance for PC1 *p =* 0.009 vs 46 (44-49)% for any pattern of PC1 variance derived from random data; Narayanan, 2016). Consistent with population averages from Fig 2G&H, D2-MSNs and D1-MSNs had opposite patterns of activity with negative PC1 scores for D2-MSNs and positive PC1 scores for D1-MSNs (Fig 3C; PC1 for D2-MSNs: -3.4 (-4.6 – 2.5); PC1 for D1-MSNs: 2.8 (-2.8 – 4.9); F = 8.8, *p* = 0.004 accounting for variance between mice (Fig S3A); Cohen’s *d* = 0.7; power = 0.80; no reliable effect of sex (F = 0.44, *p =* 0.51) or switching direction (F = 1.73, *p =* 0.19)). Importantly, PC1 scores for D2- MSNs were significantly less than 0 (signrank D2-MSN PC1 scores vs 0: *p* = 0.02), implying that because PC1 ramps down, D2-MSNs tended to ramp up. Conversely, PC1 scores for D1- MSNs were significantly greater than 0 (signrank D1-MSN PC1 scores vs 0: *p* = 0.05), implying that D1-MSNs tended to ramp down. Thus, analysis of PC1 in Fig 3A-C suggested that D2- MSNs (Fig 2G) and D1-MSNs (Fig 2H) had opposing ramping dynamics.

**Figure 3:**
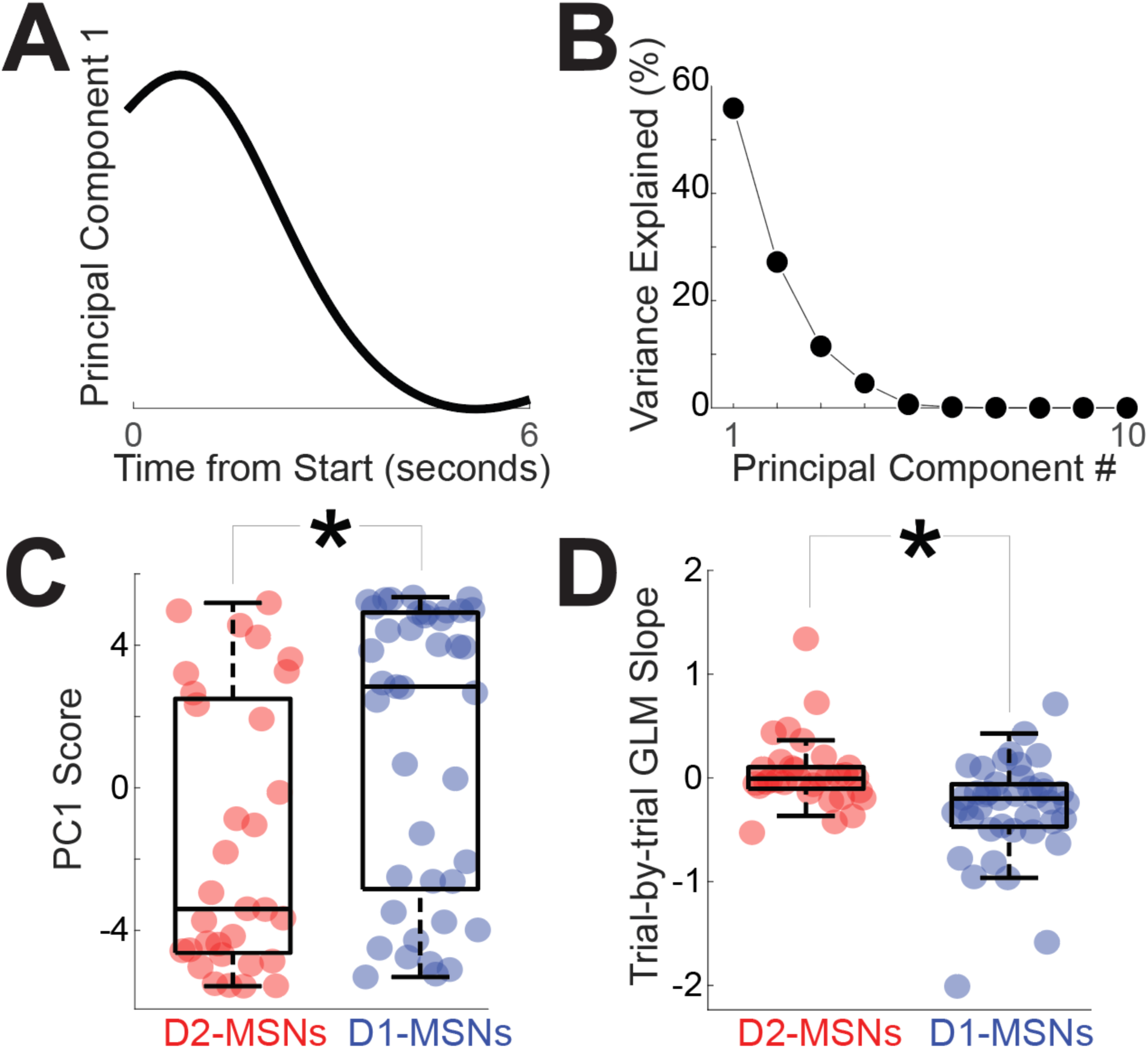
Quantification of opposing D2-MSN and D1-MSN dynamics. A) Principal component analysis revealed that the first component (PC1) exhibited time-dependent ramping. B) The first principal component explained ∼54% of variance across tagged MSN ensembles. C) Differences between D2-MSNs (red) and D1-MSNs (blue) were captured by PC1 which exhibited time-dependent ramping. D) These differences were also apparent in the linear slope of firing rate vs time in the interval, with D1-MSNs (blue) having a more negative slope than D2-MSNs (red). In C and D, each point represents data from a tagged MSN. * = *p* < 0.05 via linear mixed effects models accounting for variance between mice. Data from 32 tagged D2-MSNs in 4 D2-Cre mice and 41 tagged D1-MSNs in 5 D1-Cre mice.

To interrogate these dynamics at a trial-by-trial level, we calculated the linear slope of D2-MSN and D1-MSN activity over the first 6 seconds of each trial using generalized linear modeling (GLM) of effects of time in the interval vs trial-by-trial firing rate (Latimer et al., 2015). Note that this analysis focuses on each trial rather than population averages as in Fig 2G- H and Fig 3A-C. Nosepokes were included as a regressor for movement. GLM analysis also demonstrated that D2-MSNs had significantly different slopes (-0.01 spikes/second (-0.10 – 0.10)), which were distinct from D1-MSNs (-0.20 (-0.47– -0.06; Fig 3D; F = 8.9, *p* = 0.004 accounting for variance between mice (Fig S3B); Cohen’s *d* = 0.8; power = 0.98; no reliable effect of sex (F = 0.02, *p =* 0.88) or switching direction (F = 1.72, *p =* 0.19)). We found that D2- MSNs and D1-MSNs had significantly different slopes even when excluding outliers (4 outliers excluded outside of 95% confidence intervals; F = 7.51, *p =* 0.008 accounting for variance between mice) and when the interval was defined as the time between trial start and the switch response on a trial-by-trial basis for each neuron (F = 4.3, *p =* 0.04 accounting for variance between mice). Trial-by-trial GLM slope was correlated with PC1 scores in Fig 3A-C (PC1 scores vs GLM slope r = -0.60, *p =* 10^-8^). These data demonstrate that D2-MSNs and D1-MSNs had distinct slopes of firing rate across the interval and were consistent with analyses of average activity and PC1, which exhibited time-related ramping.

Our findings could not be easily explained by movement because 1) few responses occurred before 6 seconds (23%) and 44% of tagged neurons were recorded in trials without a single nosepoke before 6 seconds (Fig 1B-C), 2) our GLM included a regressor accounting for the nosepokes when present, and 3) nosepoke GLM βs were not reliably different between D2- MSNs and D1-MSNs (F = 1.5, *p =* 0.22 accounting for variance between mice).

In summary, we used optogenetic tagging to record from D2-MSNs and D1-MSNs during interval timing. Analyses of average activity, PC1, and trial-by-trial firing-rate slopes over the interval provide convergent evidence that D2-MSNs and D1-MSNs had distinct dynamics during interval timing. These data provide insight into temporal processing by striatal MSNs.

### Drift-diffusion models of opposing D2-MSN and D1-MSN dynamics

Our analysis of average activity (Fig 2G-H) and PC1 (Fig 3A-C) suggested that D2- MSNs and D1-MSNs might have opposing dynamics. However, past computational models of interval timing have relied on drift-diffusion dynamics that increases over the interval and accumulates evidence over time (Nguyen et al., 2020; Simen et al., 2011). To reconcile how these MSNs might complement to effect temporal control of action, we constructed a four- parameter drift-diffusion model (DDM). Our goal was to construct a DDM inspired by average differences in D2-MSNs and D1-MSNs that predicted switch-response time behavior. We constructed a DDM where *x* represents the neuronal firing rate of an ‘output’ unit collecting evidence on the activity of striatal D2-MSNs or D1-MSNs, *t* represents time measured in seconds, and *dx* and *dt* mean “change” in *x* and *t* respectively (equivalent to the derivative

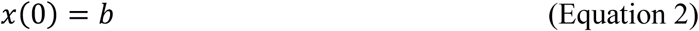

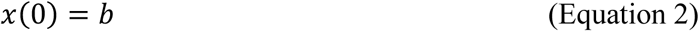

The model has four independent parameters *F*, *D*, *σ*, and *b* (described below) and a threshold value *T* defined by

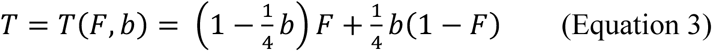

The firing rate is set initially at baseline *b* (see Equation 2), then driven by input *F* akin to the dendritic current induced by the overlap of corticostriatal postsynaptic potentials (Shepherd, 2013). With each unit of time *dt*, we suppose there is corticostriatal stimulation that provides incremental input proportional to the activity itself, (*F* − *x*)*D*, to reach a decision. *D* is *the drift rate* for the event sequence and can be interpreted as a parameter inverse proportional to the neural activity’s integration time constant. The drift, together with noise *ξ*(*t*) (of zero mean and strength *σ*), leads to fluctuating accumulation which eventually crosses a threshold *T* (see Equation 3; Fig 4A-B). The time *t*^∗^ it takes the firing rate to reach the threshold, *x*(*t*^∗^) = *T*, is the *switch response time*.

**Figure 4:**
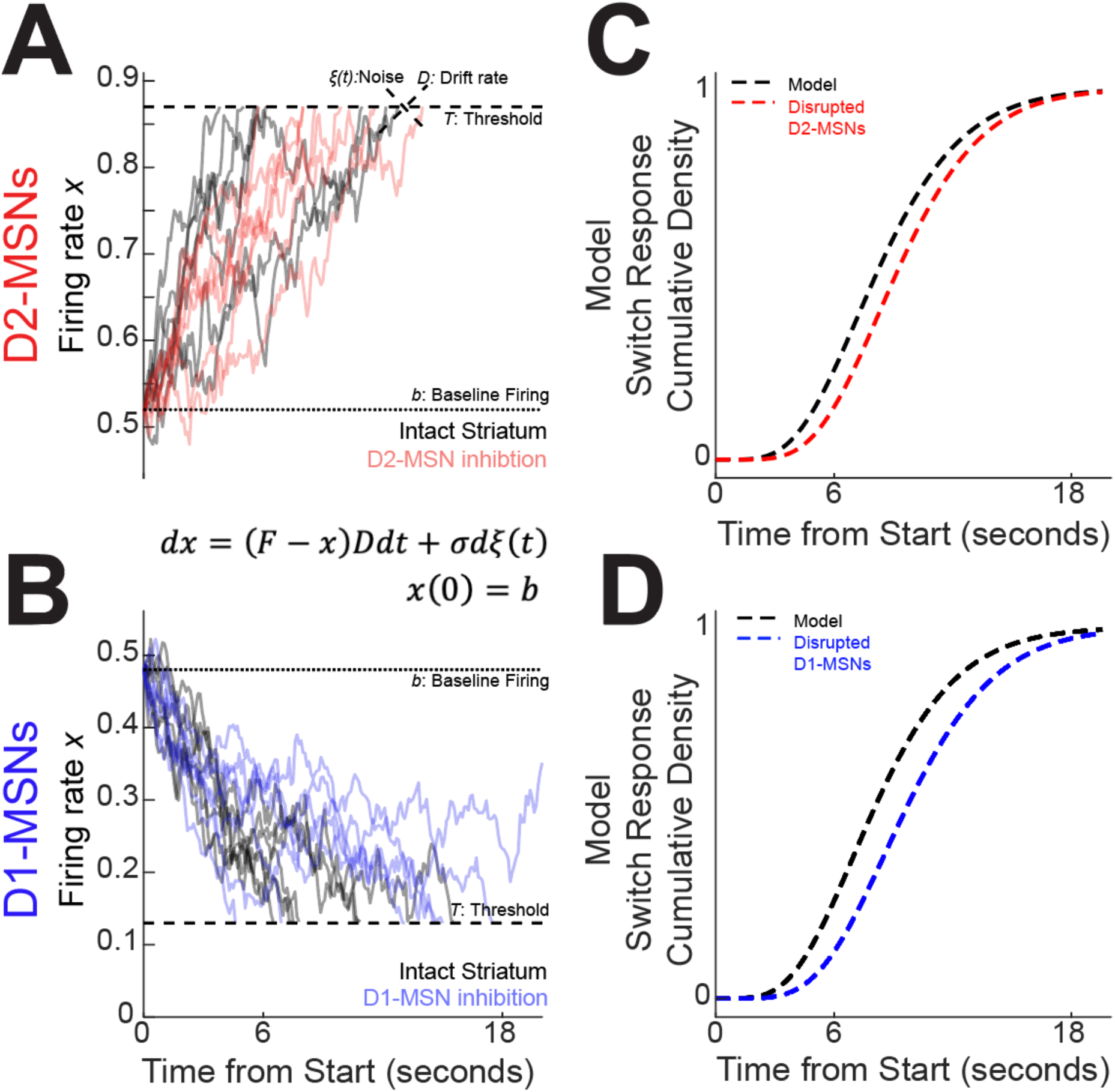
Four-parameter drift-diffusion computational model of striatal activity during interval timing. A) We modeled interval timing with a low parameter diffusion process with a drift rate *D*, noise *ξ*(*t*), and a baseline firing rate *b* that drifts toward a threshold *T* indicated by dotted lines. With D2-MSNs disrupted (solid red curves), this drift process decreases and takes longer to reach the threshold. B) The same model also accounted for D1-MSNs with an opposite drift. With D1-MSNs disrupted (solid blue curves), the drift process again takes longer to reach the threshold. Because both D2-MSNs and D1-MSNs contribute to the accumulation of temporal evidence, this model predicted that C) disrupting D2-MSNs would increase response times during interval timing (dotted red line) and D) disrupting D1-MSNs would also increase response times (dotted blue line). Threshold *T* depends on *b* and target firing *F*. For details on the selection of parameter values in DDM, see Methods and Fig S7-S8.

Our model aimed to fit statistical properties of mouse behavioral responses while incorporating MSN network dynamics. The model does not attempt, however, to fit individual neurons’ activity, because our model predicts a single behavioral parameter – switch time - that can be caused by a diversity of neuronal activity. We first analyzed trial-based aggregated activity of MSN recordings from each mouse (*x_j_*(*t*)) where *j* = 1, …, *N* neurons. For D2-MSN or D1-MSN ensembles of *N* > 11, we found linear combinations of their neuronal activities, with some *β_j_* coefficients,

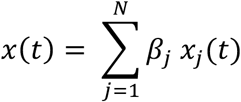

that could predict the trial-by-trial switch response times (accuracy > 90%, Fig S4; compared with < 20% accuracy for Poisson-generated spikes of same trial-average firing rate). The predicted switch time *t*^∗^*_pred_* was defined by the time when the weighted ensemble activity *x*(*t*) first reached the value *x*(*t*^∗^*_pred_*) = 0.5. Finally, we built DDMs to account for this opposing trend (increasing vs decreasing) of MSN dynamics and for ensemble threshold behavior defining *t*^∗^*_pred_*; see the resulting model (Equations 1-3) and its simulations (Figure 4A-B).

This instantiation of DDMs captured the complimentary D2-MSN and D1-MSN dynamics that we discovered in our optogenetic tagging experiments (Fig 2G-H vs 4A-B). The model’s parameters were chosen to fit the distribution of switch response times: *F* = 1, *b* = 0.52 (so *T* = 0.87), *D* = 0.135, *σ* = 0.052 for intact D2-MSNs (Fig 4A, in black); and *F* = 0, *b* = 0.48 (so *T* = 0.12), *D* = 0.141, *σ* = 0.052 for intact D1-MSNs (Fig 4B, in black). See Methods and Fig S7 for how the model parameters were chosen and how we quantified the model’s explanatory power for behavior. Interestingly, we observed that two-parameter gamma distributions strongly accounted for the model dynamics (D2-MSNs: Gamma parameters α = 6.08, β = 0.69, R^2^ Gamma vs Model = 0.99; D1-MSNs α = 5.89, β = 0.69; R^2^ Gamma vs Model = 0.99; Fig 4C-D, dotted black lines). Gamma distributions also provided a surprisingly good approximation for the probability distribution and cumulative distribution functions of mouse switch response times (Fig 1B-D; R^2^ Data vs Gamma =0.94 for D1-MSNs and for D2-MSNs). Thus, in the next sections, we used gamma distributions as a proxy for both the distribution of times generated by the model and the distribution of mice switch response times, when comparing those for the goodness of fit. Our model provided the opportunity to computationally explore the consequences of disrupting D2-MSNs or D1-MSNs. Because both D2-MSNs and D1-MSNs accumulate temporal evidence, disrupting either MSN type in the model changed the slope. The results were obtained by simultaneously decreasing the drift rate *D* (equivalent to lengthening the neurons’ integration time constant) and lowering the level of network noise *σ*: *D* = 0.129, *σ* = 0.043 for D2-MSNs in Fig 4A (in red; changes in noise had to accompany changes in drift rate to preserve switch response time variance. See Methods); and *D* = 0.122, *σ* = 0.043 for D1-MSNs in Fig 4B (in blue). The model predicted that disrupting either D2-MSNs or D1-MSNs would increase switch response times (Fig 4C and Fig 4D) and would shift MSN dynamics. In the next section, we interrogated these ideas with a combination of optogenetics, behavioral pharmacology, and electrophysiology.

### Disrupting D2-MSNs or D1-MSNs increases switch response times

DDMs captured opposing MSN dynamics and predicted that disrupting either D2-MSNs or D1-MSNs should slow temporal processing and increase switch response times (Fig 4). We tested this idea with optogenetics. We bilaterally implanted fiber optics and virally expressed the inhibitory opsin halorhodopsin in the dorsomedial striatum of 10 D2-Cre mice (5 female) to inhibit D2-MSNs (Fig S5; this group of mice was entirely separate from the optogenetic tagging mice). We found that D2-MSN inhibition reliably increased switch response times (Fig 5A-B; Laser Off: 8.6 seconds (8.3–9.3); Laser On: 10.2 seconds (9.4–10.2); signed rank *p* = 0.002, Cohen’s *d* = 1.7). To control for heating and nonspecific effects of optogenetics, we performed control experiments in D2-cre mice without opsins using identical laser parameters; we found no reliable effects for opsin-negative controls (Fig S6). Remarkably, DDM predictions were highly concordant with D2-MSN inhibition behavioral data (R^2^ Data vs Model= 0.95; Fig S7-S8) as well as with behavioral data from laser off trials (R^2^ Data vs Model= 0.94; parameters chosen to fit laser off trials; Fig S7).

**Figure 5:**
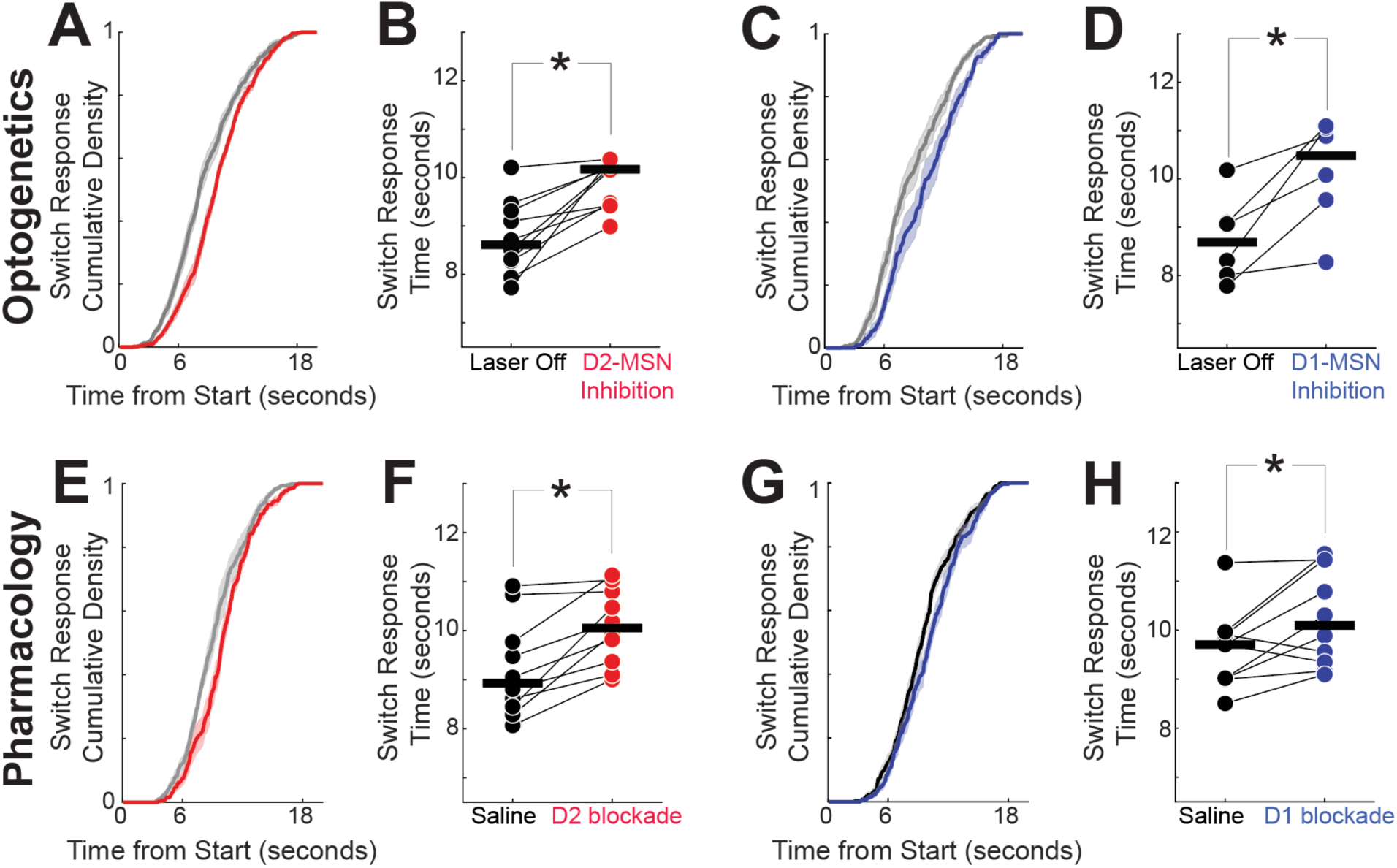
Disrupting D2- or D1-MSNs increases response times. A) As predicted by our DDM in Fig 4, optogenetic inhibition of D2-MSNs (red) shifted cumulative distributions of response times to the right, and B) increased response times; data from 10 D2-Cre mice expressing halorhodopsin (Halo). Also as predicted by our DDM, C) optogenetic inhibition of D1-MSNs shifted cumulative distribution functions to the right, and D) increased response times; data from 6 D1-Cre mice expressing Halo. Similarly, E) pharmacologically disrupting D2-dopamine receptors (red) with the D2 antagonist sulpiride shifted cumulative distribution functions to the right, and F) increased response times; data from 10 wild-type mice. Also, G) pharmacologically disrupting D1-dopamine receptors (blue) with the D1 antagonist SCH23390 shifted cumulative distribution functions to the right, and H) increased response times; data from the same 10 wild-type mice as in E-F. In B, D, F, and H connected points represent the mean response time from each animal in each session, and horizontal black lines represent group medians. \**p* = < 0.05, signed rank test. See Figure S6 for data from opsin- negative controls.

Next, we investigated D1-MSNs (Kravitz et al., 2010). In 6 D1-Cre mice (3 female), optogenetic inhibition of dorsomedial striatal D1-MSNs increased switch response times (Fig 5C-D; Laser Off: 8.7 (8.0–9.1) seconds; Laser On: 10.5 (9.6 – 11.1) seconds; signed rank *p* = 0.03, Cohen’s *d* = 1.4). As with D2-MSNs, we found no reliable effects with opsin-negative controls in D1-MSNs (Fig S6). DDM predictions again were highly concordant with D1-MSN inhibition behavioral data (R^2^ Data vs Model= 0.95; parameters chosen to fit D1-MSN inhibition trials and with laser off trials (R^2^ Data vs Model= 0.94; Fig S7-S8).

To test the generality of the effects observed in the optogenetic inhibition experiment, we turned to systemic pharmacology. We used sulpiride (12.5 mg/kg intraperitoneally (IP)) to block D2-dopamine receptors in 10 wild-type (WT) mice, which increased switch response times relative to saline sessions (median (IQR); saline: 8.9 (8.4 – 9.8) seconds; D2 blockade: 10.1 (9.4 – 10.8) seconds; signed rank *p* = 0.002, Cohen’s *d* = 1.0; Fig 5E-F). There was no difference in switch response time between saline sessions and laser-off trials from D2-MSN optogenetic inhibition in D2-Cre mice, implying that in optogenetic sessions, D2-MSN inhibition did not disrupt behavior during laser-off trials (rank sum *p* = 0.31). These data are consistent with past work demonstrating that disrupting dorsomedial D2-MSNs slows timing (De Corte et al., 2019b; Drew et al., 2007; Stutt et al., 2024). In the same 10 wild-type mice, systemic drugs blocking D1- dopamine receptors (D1 blockade; SCH23390 0.05 mg/kg IP) increased switch response times (saline: 9.7 seconds (9.0 – 9.9); D1 blockade: 10.1 seconds (9.3 – 11.4); signed rank *p* = 0.04, Cohen *d* = 0.7; Fig 5G-H). Once again, there was no difference between saline sessions and D1- MSN inhibition laser-off trials in D1-Cre mice (rank sum *p* = 0.18). These results are in agreement with our DDM predictions and demonstrate that disrupting either D2-MSNs or D1- MSNs increases switch response time.

We found no evidence that inhibiting D2-MSNs in the dorsomedial striatum changed task-specific movements such as nosepoke duration (i.e., time of nosepoke entry to exit; signed rank *p* = 0.63; Fig S9 for the 10 D2-Cre mice in optogenetics trials) or switch traversal time between the first and second nosepokes (signed rank *p* = 0.49; Fig S9). Similarly, we found no evidence that D1-MSN inhibition changed nosepoke duration (signed rank *p* = 0.31; Fig S9 for the 6 D1-Cre mice) or traversal time (signed rank *p* = 0.22; Fig S9). Furthermore, disrupting D2- MSNs or D1-MSNs did not change switch response time standard deviations or the number of rewards (Fig S10). These results demonstrate that disrupting dorsomedial striatal D2-MSNs and D1-MSNs specifically slowed interval timing without consistently changing task-specific movements.

### D2 blockade and D1 blockade shift MSN dynamics

MSN ensembles strongly encode time (Bruce et al., 2021; Emmons et al., 2017; Gouvea et al., 2015; Mello et al., 2015; Wang et al., 2018), but it is unknown how disruptions in D2- dopamine or D1-dopamine receptors affect these ensembles (Yun et al., 2023). Although non- specific, pharmacological experiments have two advantages over optogenetics for recording experiments: 1) many clinically approved drugs target dopamine receptors, and 2) they are more readily combined with recordings than optogenetic inhibition, which silences large populations of MSNs. We recorded from dorsomedial striatal MSN ensembles in 11 separate mice during sessions with saline, D2 blockade with sulpiride, or D1 blockade with SCH23390 (Fig 6A; Fig S11; data from one recording session each for saline, D2 blockade, and D1 blockade during interval timing).

**Figure 6:**
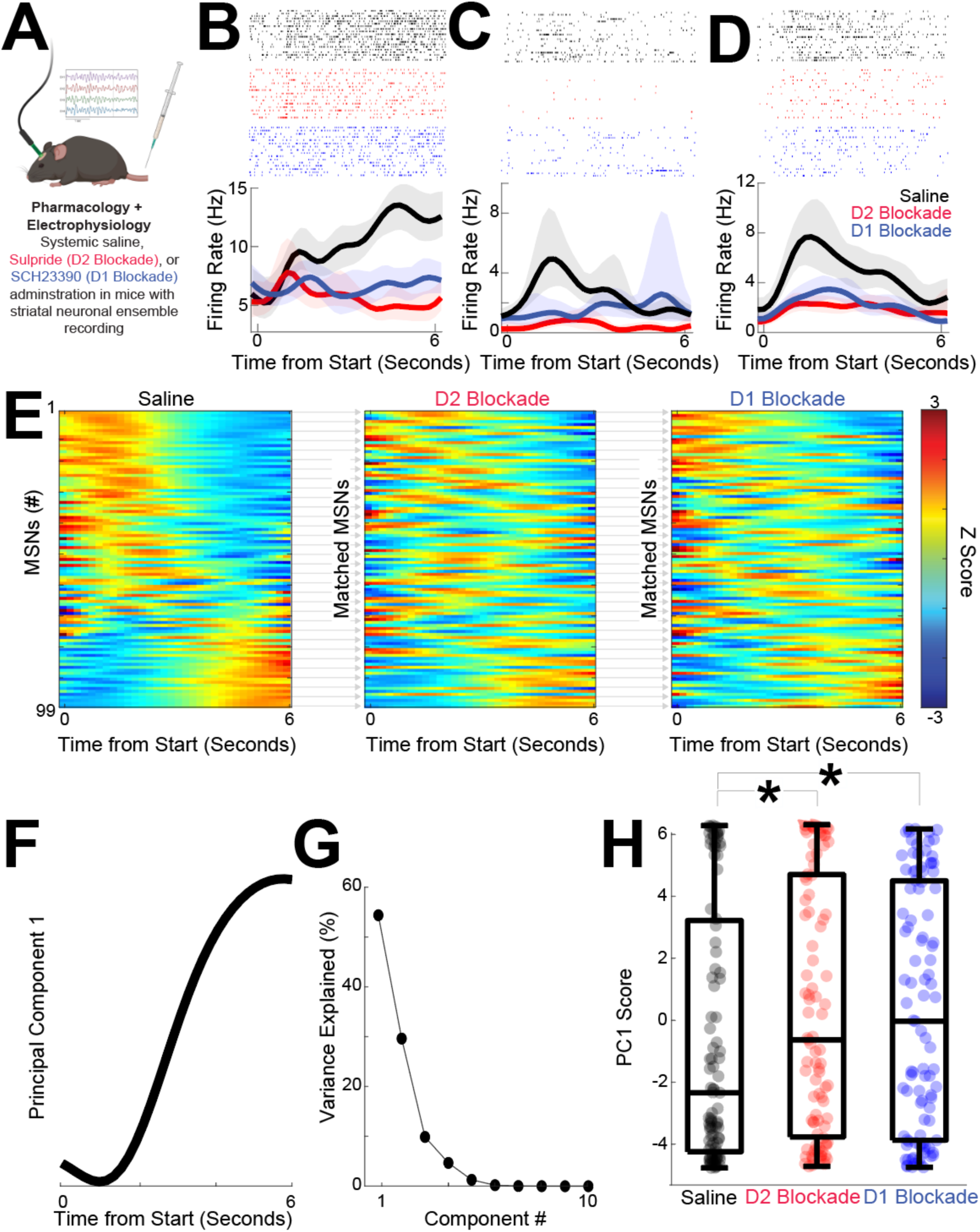
D2 and D1 blockade shift temporal dynamics. A) We recorded dorsomedial striatal medium spiny neuron (MSN) ensembles during interval timing in sessions with saline, D2 blockade with sulpiride, or D1 blockade with SCH23390. Made with BioRender.com. B-D) Example peri-event raster from MSNs in sessions with saline (black), D2- dopamine blockade (red), or D1-dopamine blockade (blue). Shaded area is the bootstrapped 95% confidence interval. E) MSNs from 99 neurons in 11 mice from saline, D2 blockade, or D1 blockade session; MSNs were matched across sessions based on waveforms and interspike interval. Each row represents a peri-event time histogram (PETH) binned at 0.2 seconds, smoothed using kernel-density estimates using a bandwidth of 1, and z-scored. Colors indicate z-scored firing rate. See Fig S11 for analyses that assume statistical independence. F) Principal component analysis (PCA) identified MSN ensemble patterns of activity. The first principal component (PC1) exhibited time-dependent ramping. G) PC1 explained 54% of population variance among MSN ensembles; higher components were not analyzed. H) PC1 scores were closer to zero and significantly different with D2 or D1 blockade; * = *p* < 0.05 via linear mixed effects; data from 99 MSNs in 11 mice.

We analyzed 99 MSNs in sessions with saline, D2 blockade, and D1 blockade. We matched MSNs across sessions based on waveform and interspike intervals; waveforms were highly similar across sessions (correlation coefficient between matched MSN waveforms: saline vs D2 blockade r = 1.00 (0.99 – 1.00 rank sum vs correlations in unmatched waveforms *p* = 3x10^-44;^ waveforms; saline vs D1 blockade r = 1.00 (1.00 – 1.00), rank sum vs correlations in unmatched waveforms *p* = 4x10^-50^). There were no consistent changes in MSN average firing rate with D2 blockade or D1 blockade (F = 1.1, *p =* 0.30 accounting for variance between MSNs; saline: 5.2 (3.3 – 8.6) Hz; D2 blockade 5.1 (2.7 – 8.0) Hz; F = 2.2, *p* = 0.14; D1 blockade 4.9 (2.4 – 7.8) Hz).

We noticed differences in MSN activity across the interval with D2 blockade and D1 blockade at the individual MSN level (Fig 6B-D) as well as at the population level (Fig 6E). We used PCA to quantify effects of D2 blockade or D1 blockade (Bruce et al., 2021; Emmons et al., 2017; Kim et al., 2017a). We constructed principal components (PC) from z-scored peri-event time histograms of firing rate from saline, D2 blockade, and D1 blockade sessions for all mice together. The first component (PC1), which explained 54% of neuronal variance, exhibited “time-dependent ramping”, or monotonic changes over the 6 second interval immediately after trial start (Fig 6F-G; variance for PC1 *p* = 0.001 vs 46 (45-47)% for any pattern of PC1 variance in random data; Narayanan, 2016). Interestingly, PC1 scores shifted with D2 blockade (Fig 6F; PC1 scores for D2 blockade: -0.6 (-3.8 – 4.7) vs saline: -2.3 (-4.2 – 3.2), F = 5.1, *p* = 0.03 accounting for variance between MSNs; no reliable effect of sex (F = 0.2, *p* = 0.63) or switching direction (F = 2.8, *p* = 0.10)). PC1 scores also shifted with D1 blockade (Fig 6F; PC1 scores for D1 blockade: -0.0 (-3.9 – 4.5), F = 5.8, *p* = 0.02 accounting for variance between MSNs; no reliable effect of sex (F = 0.0, *p* = 0.93) or switching direction (F = 0.9, *p* = 0.34)). There were no reliable differences in PC1 scores between D2 and D1 blockade. Furthermore, PC1 was distinct even when sessions were sorted independently and assumed to be fully statistically independent (Figure S11; D2 blockade vs saline: F = 5.5, *p =* 0.02; D1 blockade vs saline: F = 4.9, *p =* 0.03; all analyses accounting for variance between mice). Higher components explained less variance and were not reliably different between saline and D2 blockade or D1 blockade. Taken together, this data-driven analysis shows that D2 and D1 blockade produced similar shifts in MSN population dynamics represented by PC1. When combined with the major contributions of D1/D2 MSNs to PC1 (Fig 3C) these findings indicate that pharmacological D2 blockade and D1 blockade disrupt ramping-related activity in the striatum.

### D2 blockade and D1 blockade degrades MSN temporal encoding

Finally, we quantified striatal MSN temporal encoding via a naïve Bayesian classifier that generates trial-by-trial predictions of time from MSN ensemble firing rates (Fig 7A-C; (Bruce et al., 2021; Emmons et al., 2017). Our DDMs predict that disrupted temporal decoding would be a consequence of an altered DDM drift rate. We used leave-one-out cross-validation to predict objective time from the firing rate within a trial. Saline sessions generated strong temporal predictions for the first 6 seconds of the interval immediately after trial start (0–6 seconds; R^2^ = 0.91 (0.83 – 0.94)) with weaker predictions for later epochs (6–12 seconds: R^2^ = 0.55 (0.34 – 0.70); rank sum *p* = 0.00002 vs 0–6 seconds, Cohen *d* = 2.0; 12–18 seconds: R^2^ = 0.18 (0.10 – 0.62); rank sum *p* = 0.00003 vs 0–6 seconds, Cohen *d* = 2.4; all analyses considered each epoch statistically independent; Fig 7D). We found that temporal encoding early in the interval (0–6 seconds) was degraded with either D2 blockade (R^2^ = 0.69 (0.58 – 0.84); rank sum *p* = 0.0002 vs saline, Cohen *d* = 1.4) or D1 blockade (R^2^ = 0.71 (0.47 – 0.87); rank sum *p* = 0.004 vs saline, Cohen *d* = 1.1; Fig 7D), consistent with predictions made from DDMs. Later in the interval (6–12 and 12-18 seconds), there were no significant differences between saline sessions and D2 blockade or D1 blockade (Fig 7D).

**Figure 7:**
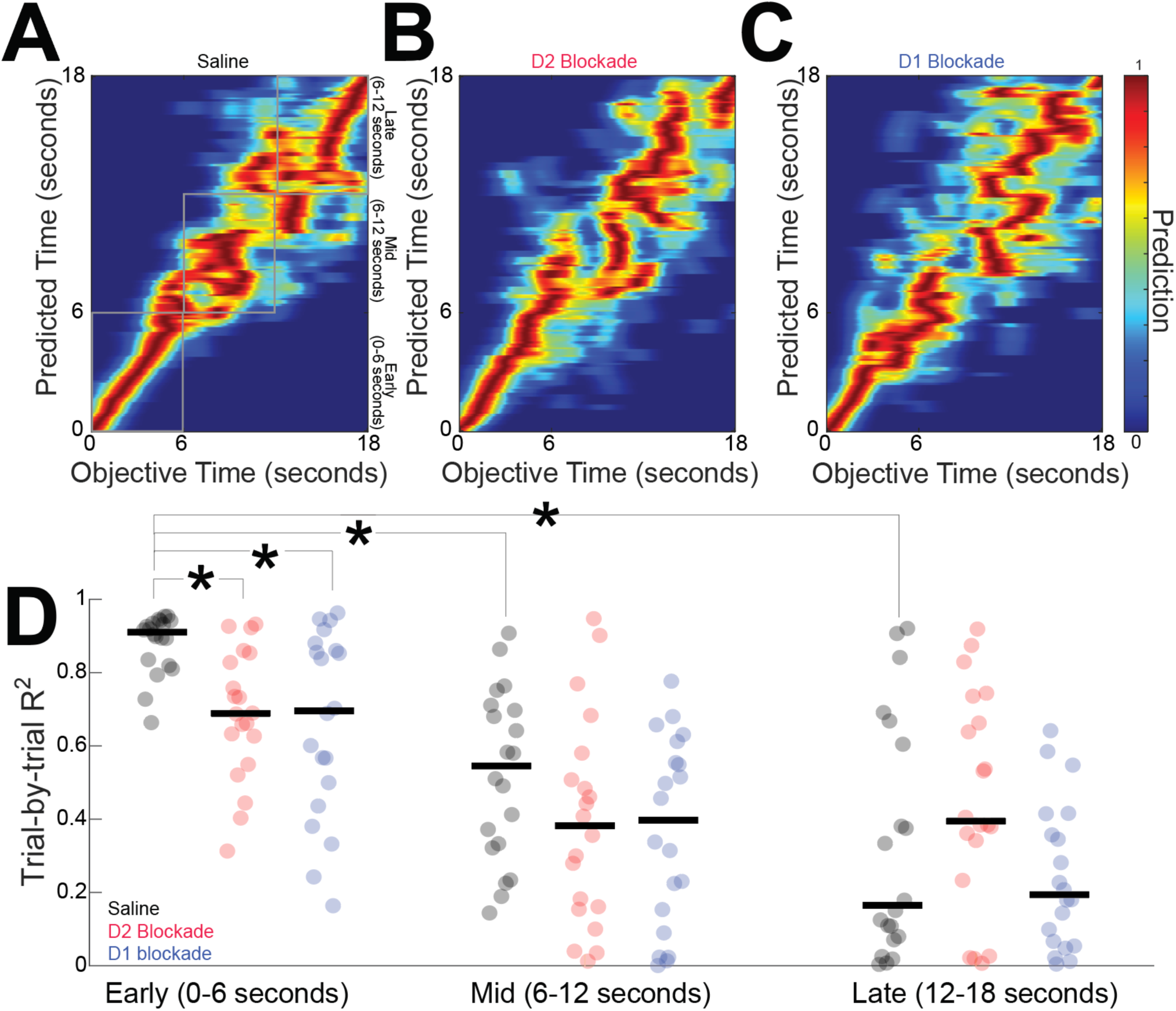
D2 and D1 blockade degrade MSN temporal encoding. We used naïve Bayesian classifiers to decode time from MSN ensembles in A) saline sessions, B) D2 blockade sessions, and C) D1 blockade sessions. Color represents the temporal prediction across 20 trials with red representing stronger predictions. D) Temporal encoding was strong early in the interval, and D2 or D1 blockade degraded classification accuracy. Temporal encoding was decreased later in the interval. Each point represents the R^2^ for each trial of behavior for MSN ensembles from 11 mice. * *p* < 0.05 vs saline from 0-6 seconds. Horizontal black lines in (D) represent group medians.

Taken together, these data demonstrate that disrupting either D2-MSNs or D1-MSNs degrades temporal encoding by MSN ensembles. In combination with our optogenetic tagging, computational modeling, and optogenetic inhibition experiments, these data provide insight into cognitive computations by the striatum.

## DISCUSSION

We describe how striatal MSNs work together in complementary ways to encode an elementary cognitive process, interval timing. Strikingly, optogenetic tagging showed that D2- MSNs and D1-MSNs had distinct dynamics during interval timing. MSN dynamics helped construct and constrain a four-parameter drift-diffusion model in which D2- and D1-MSN spiking accumulated temporal evidence. This model predicted that disrupting either D2-MSNs or D1-MSNs would increase switch response times. Accordingly, we found that optogenetically or pharmacologically disrupting striatal D2-MSNs or D1-MSNs increased switch response times without affecting task-specific movements. Disrupting D2-MSNs or D1-MSNs shifted MSN temporal dynamics and degraded MSN temporal encoding. These data, when combined with our model predictions, demonstrate that despite opposing dynamics, D2-MSNs and D1-MSN contribute complementary temporal evidence to controlling actions in time. Our interpretation is that because the activity of D2-MSN and D1-MSN ensembles represents the accumulation evidence, pharmacological/optogenetic disruption of D2-MSN/D1-MSN activity slows this accumulation process, leading to slower interval timing-response times (Fig 5) without changing other task-specific movements (Fig S9). These results provide new insight into how opposing patterns of striatal MSN activity control behavior in similar ways and show that they play a complementary role in elementary cognitive operations.

Striatal MSNs are critical for temporal control of action (Emmons et al., 2017; Gouvea et al., 2015; Mello et al., 2015). Three broad models have been proposed for how striatal MSN ensembles represent time: 1) the striatal beat frequency model, in which MSNs encode temporal information based on neuronal synchrony (Matell and Meck, 2004); 2) the distributed coding model, in which time is represented by the state of the network (Paton and Buonomano, 2018); and 3) the DDM, in which neuronal activity monotonically drifts toward a threshold after which responses are initiated (Emmons et al., 2017; Simen et al., 2011; Wang et al., 2018). While our data do not formally resolve these possibilities, our results show that D2-MSNs and D1-MSNs exhibit opposing changes in firing rate dynamics in PC1 over the interval. Past work by our group and others has demonstrated that PC1 dynamics can scale over multiple intervals to represent time (Emmons et al., 2020, 2017; Gouvea et al., 2015; Mello et al., 2015; Wang et al., 2018). We find that low-parameter DDMs account for interval timing behavior with both intact and disrupted striatal D2- and D1-MSNs. While other models can capture interval timing behavior and account for MSN neuronal activity, our model does so parsimoniously with relatively few parameters (Matell and Meck, 2004; Paton and Buonomano, 2018; Simen et al., 2011). We and others have shown previously that ramping activity scales to multiple intervals, and DDMs can be readily adapted by changing the drift rate (Emmons et al., 2017; Gouvea et al., 2015; Mello et al., 2015; Simen et al., 2011). Interestingly, decoding performance was high early in the interval; indeed, animals may have been focused on this initial interval (Balci and Gallistel, 2006) in making temporal comparisons and deciding whether to switch response nosepokes.

D2-MSNs and D1-MSNs play complementary roles in movement. For instance, stimulating D1-MSNs facilitates movement, whereas stimulating D2MSNs impairs movement (Kravitz et al., 2010). Both populations have been shown to have complementary patterns of activity during movements with MSNs firing at different phases of action initiation and selection (Tecuapetla et al., 2016). Further dissection of action selection programs reveals that opposing patterns of activation among D2-MSNs and D1-MSNs suppress and guide actions, respectively, in the dorsolateral striatum (Cruz et al., 2022). A particular advantage of interval timing is that it captures a cognitive behavior within a single dimension — time. When projected along the temporal dimension, it was surprising that D2-MSNs and D1-MSNs had opposing patterns of activity. Ramping activity in the prefrontal cortex can increase or decrease; and prefrontal neurons project to and control striatal ramping activity (Emmons et al., 2020, 2017; Wang et al., 2018). It is possible that differences in D2-MSNs and D1-MSNs reflect differences in cortical ramping, which may themselves reflect more complex integrative or accumulatory processes.

Further experiments are required to investigate these differences. Past pharmacological work from our group and others has shown that disrupting D2- or D1-MSNs slows timing (De Corte et al., 2019b; Drew et al., 2007, 2003; Stutt et al., 2024) and are in agreement with pharmacological and optogenetic results in this manuscript. Computational modeling predicted that disrupting either D2-MSNs or D1-MSNs increased self-reported estimates of time, which was supported by both optogenetic and pharmacological experiments.

Notably, these disruptions are distinct from increased timing variability reported with administrations of amphetamine, ventral tegmental area dopamine neuron lesions, and rodent models of aging (Balci et al., 2008; Gür et al., 2020; Weber et al., 2023). Furthermore, our current data demonstrate that disrupting either D2-MSN or D1-MSN activity shifted MSN dynamics and degraded temporal encoding, supporting prior work (De Corte et al., 2019b; Drew et al., 2007, 2003; Stutt et al., 2024). Our recording experiments do not identify where a possible response threshold *T* is instantiated, but downstream basal ganglia structures may have a key role in setting response thresholds (Toda et al., 2017).

Because interval timing is reliably disrupted in human diseases of the striatum such as Huntington’s disease, Parkinson’s disease, and schizophrenia (Hinton et al., 2007; Singh et al., 2021; Ward et al., 2011), these results have relevance to human disease. Our task version has been used extensively to study interval timing in mice and humans (Balci et al., 2008; Bruce et al., 2021; Stutt et al., 2024; Tosun et al., 2016; Weber et al., 2023). However, temporal bisection tasks, in which animals hold during a temporal cue and respond at different locations depending on cue length, have advantages in studying how animals time an interval because animals are not moving while estimating cue duration (Paton and Buonomano, 2018; Robbe, 2023; Soares et al., 2016). Our interval timing task version – in which mice switch between two response nosepokes to indicate their interval estimate has elapsed – has been used extensively in rodent models of neurodegenerative disease (Larson et al., 2022; Weber et al., 2024, 2023; Zhang et al., 2021), as well as in humans (Stutt et al., 2024). This version of interval timing involves motor timing, which engages executive function and has more translational relevance for human diseases than perceptual timing or bisection tasks (Brown, 2006; Farajzadeh and Sanayei, 2024; Nombela et al., 2016; Singh et al., 2021). Furthermore, because many therapeutics targeting dopamine receptors are used clinically, these findings help describe how dopaminergic drugs might affect cognitive function and dysfunction. Future studies of D2-MSNs and D1-MSNs in temporal bisection and other timing tasks may further clarify the relative roles of D2- and D1-MSNs in interval timing and time estimation.

Our approach has several limitations. First, systemic drug injections block D2- and D1- receptors in many different brain regions, including the frontal cortex, which is involved in interval timing (Kim et al., 2017a). D2 blockade or D1 blockade may have complex effects, including corticostriatal or network effects that contribute to changes in D2-MSN or D1-MSN ensemble activity. We note that optogenetic inhibition of D2-MSNs and D1-MSNs produces similar effects to pharmacology in Fig 5. Pharmacology is compatible with neuronal ensemble recordings as optogenetic inhibition would silence a large fraction of neurons, complicating interpretations. Future studies might extend our work by combining local pharmacology with neuronal ensemble recording. Second, although we had adequate statistical power and medium- to-large effect sizes, optogenetic tagging is low-yield, and it is possible that recording more of these neurons would afford greater opportunity to identify more robust results and alternative coding schemes, such as neuronal synchrony. Furthermore, we find that recording more neurons simultaneously would also help further constrain DDMs to predict trial-by-trial firing rate and generate a more sophisticated neuronal network model of time (Fig S4). Third, the striatum includes diverse cell types, some of which express both D1 and D2 dopamine receptors; these cell types may also contribute to cognitive processing. Fourth, MSNs can laterally inhibit each other, which may profoundly affect striatal function. Regardless, we show that cell-type-specific disruption of D2-MSNs and D1-MSNs both slow timing, implying that these cell types play a key role in interval timing. Future experiments may record from non-MSN striatal cell types, including fast-spiking interneurons that shape basal ganglia output. Fifth, we did not deliver stimulation to the striatum because our pilot experiments triggered movement artifacts or task- specific dyskinesias (Kravitz et al., 2010). Future stimulation approaches carefully titrated to striatal physiology may affect interval timing without affecting movement. Finally, movement and motivation contribute to MSN dynamics (Robbe, 2023). Four lines of evidence argue that our findings cannot be directly explained by motor confounds: 1) D2-MSNs and D1-MSNs diverge between 0-6 seconds after trial start well before the first nosepoke (Fig S2), 2) our GLM accounted for nosepokes and nosepoke-related βs were similar between D2-MSNs and D1- MSNs, 3) optogenetic disruption of dorsomedial D2-MSNs and D1-MSNs did not change task- specific movements despite reliable changes in switch response time, and 4) ramping dynamics were quite distinct from movement dynamics. Furthermore, disrupting D2-MSNs and D1-MSNs did not change the number of rewards animals received, implying that these disruptions did not grossly affect motivation. Still, future work combining motion tracking/accelerometry with neuronal ensemble recording and optogenetics and including bisection tasks may further unravel timing vs movement in MSN dynamics (Robbe, 2023; Tecuapetla et al., 2016).

In summary, we examined the role of dorsomedial striatal D2-MSNs and D1-MSNs during an elementary cognitive behavior, interval timing. Optogenetic tagging revealed that D2- MSNs and D1-MSNs exhibited opposite and complementary patterns of neuronal activity. These dynamics could be captured by computational drift-diffusion models, which predicted that disrupting either D2-MSNs or D1-MSNs would slow the accumulation of temporal evidence and increase switch response time. In concordance with this prediction, we found that optogenetic or pharmacological disruption of either D2-MSNs or D1-MSNs increased switch response times, with pharmacological D2 or D1 blockade shifting MSN dynamics and degrading temporal decoding. Collectively, our data provide insight into how the striatum encodes cognitive information, which could be highly relevant for human diseases that disrupt the striatum and for next-generation neuromodulation that targets the basal ganglia to avoid cognitive side effects or to treat cognitive dysfunction.

## Methods and Materials

### Rodents

All procedures were approved by the Institutional Animal Care and Use Committee (IACUC) at the University of Iowa, and all experimental methods were performed in accordance with applicable guidelines and regulations (Protocol #0062039). We used five cohorts of mice, summarized in Table 1: 1) 30 wild-type C57BL/6J mice (17 female) for behavioral experiments (Fig 1), 2) 4 *Drd2-cre*+ mice derived from Gensat strain ER44 (2 female) and 5 *Drd1-cre+* mice derived from Gensat strain EY262 (2 female) for optogenetic tagging and neuronal ensemble recordings in Fig 2-3; 3) 10 *Drd2-cre*+ (5 female), and 6 *Drd1-cre+* mice (3 female) for optogenetic inhibition (Fig 5) with 5 *Drd2-cre+* and 5 *Drd1-cre+* controls and 4) 10 wild-type mice for behavioral pharmacology (Fig 5), and 5) 11 mice (4 C57BL/6J (2 female), 5 *Drd2-cre+* mice (2 female), and 2 *Drd1-cre+* mice (0 female)) for combined behavioral pharmacology and neuronal ensemble recording (Fig 6-7) (Table 1). Our recent work shows that D2-blockade and D1-blockade have similar effects in both sexes (Stutt et al., 2024).

### Interval timing switch task

We used a mouse-optimized operant interval timing task described in detail previously (Balci et al., 2008; Bruce et al., 2021; Tosun et al., 2016; Weber et al., 2023). Briefly, mice were trained in sound-attenuating operant chambers, with two front nosepokes flanking either side of a food hopper on the front wall, and a third nosepoke located at the center of the back wall. The chamber was positioned below an 8-kHz, 72-dB speaker (Fig 1A; MedAssociates, St. Albans, VT). Mice were 85% food restricted and motivated with 20 mg sucrose pellets (Bio-Serv, Flemington, NJ). Mice were initially trained to receive rewards during fixed ratio nosepoke response trials. Nosepoke entry and exit were captured by infrared beams.

After shaping, mice were trained in the “switch” interval timing task. Mice self-initiated trials at the back nosepoke, after which tone and nosepoke lights were illuminated simultaneously. Cues were identical on all trial types and lasted the entire duration of the trial (6 or 18 seconds). On 50% of trials, mice were rewarded for a nosepoke after 6 seconds at the designated first ‘front’ nosepoke; these trials were not analyzed. On the remaining 50% of trials, mice were rewarded for nosepoking first at the ‘first’ nosepoke location and then switching to the ‘second’ nosepoke location; the reward was delivered for initial nosepokes at the second nosepoke location after 18 seconds when preceded by a nosepoke at the first nosepoke location. Multiple nosepokes at each nosepokes were allowed. Early responses at the first or second nosepoke were not reinforced.

Initial responses at the second nosepoke rather than the first nosepoke, alternating between nosepokes, going back to the first nosepoke after the second nosepoke were rare after initial training. Error trials included trials where animals responded only at the first or second nosepoke and were also not reinforced. We did not analyze error trials as they were often too few to analyze; these were analyzed at length in our prior work (Bruce et al., 2021).

*Switch response time* was defined as the moment animals departed the first nosepoke before arriving at the second nosepoke. Critically, switch responses are a time-based decision guided by temporal control of action because mice switch nosepokes only if nosepokes at the first location did not receive a reward after 6 seconds. That is, mice estimate if more than 6 seconds have elapsed without receiving a reward to decide to switch responses. Mice learn this task quickly (3-4 weeks), and error trials in which an animal nosepokes in the wrong order or does not nosepoke are relatively rare and discarded. Consequently, we focused on these switch response times as the key metric for temporal control of action. *Traversal time* was defined as the duration between first nosepoke exit and second nosepoke entry and is distinct from switch response time when animals departed the first nosepoke. *Nosepoke duration* was defined as the time between first nosepoke entry and exit for the switch response times only. Trials were self-initiated, but there was an intertrial interval with a geometric mean of 30 seconds between trials.

### Surgical and histological procedures

Surgical procedures were identical to methods described previously (Bruce et al., 2021). Briefly, mice were anesthetized using inhaled 4% isoflurane and surgical levels of anesthesia were maintained at 1-2% for the duration of the surgery.

Craniotomies were drilled above bilateral dorsal striatal anatomical targets, and optogenetic viruses (AAV5-DIO-eNHPR2.0 (halorhodopsin), AAV5-DIO-ChR2(H134R)-mcherry (ChR2), or AAV5-DIO-cherry (control) from the University of North Carolina Viral Vector Core) were injected into the dorsal striatum (0.5 uL of virus, +0.9, ML +/-1.3, DV –2.7). Either fiber optics (Doric Lenses, Montreal Quebec; AP +0.9, ML +/-1.3, DV –2.5) or 4 x 4 electrode or optrode arrays (AP +0.4, ML -1.4, DV -2.7 on the left side only; Microprobes, Gaithersburg, MD) were positioned in the dorsal striatum. Holes were drilled to insert skull screws to anchor headcap assemblies and/or ground electrode arrays and then sealed with cyanoacrylate (“SloZap,” Pacer Technologies, Rancho Cucamonga, CA), accelerated by “ZipKicker” (Pacer Technologies) and methyl methacrylate (AM Systems, Port Angeles, WA). Following postoperative recovery, mice were trained on the switch task and acclimated to experimental procedures prior to undergoing experimental sessions.

### Optogenetics

We leveraged cell-type-specific optogenetics to manipulate D2-MSNs and D1- MSNs in D2-cre or D1-cre mice. In animals injected with optogenetic viruses, optical inhibition was delivered via bilateral patch cables for the entire trial duration of 18 seconds via 589-nm laser light at 12 mW power on 50% of randomly assigned trials. To control for heating and nonspecific effects of optogenetics, we performed control experiments in mice without opsins using identical laser parameters in D2-cre or D1-cre mice (Fig S6). We did not stimulate for epochs less than the interval because we did not want to introduce a cue during the interval. For optogenetic tagging, putative D1- and D2-MSNs were optically identified via 473-nm photostimulation. Units with mean post-stimulation spike latencies of ≤5 milliseconds and a stimulated-to-unstimulated waveform correlation ratio of >0.9 were classified as putative D2- MSNs or D1-MSNs (Ryan et al., 2018; Shin et al., 2018). Only one recording session was performed for each animal per day, and one recording session was included from each animal.

### Behavioral pharmacology procedures

C57BL/6J mice were injected intraperitoneally (IP) 20-40 minutes before interval timing trials with either SCH23390 (C17H18ClNO; D1 antagonist), sulpiride (C15H23N3O4S; D2 antagonist), or isotonic saline. The sulpiride dosage was 12.5 mg/kg, 0.01 mL/g, and the SCH23390 was administered at a dosage of 0.05 mg/kg, 0.01 mL/g.

Behavioral performance was compared with interval timing behavior on the prior day when isotonic saline was injected IP. Only one recording session was performed for each animal per day, and one recording session was included from saline, D2 blockade, and D1 blockade sessions.

### Electrophysiology

Single-unit recordings were made using a multi-electrode recording system (Open Ephys, Atlanta, GA). After the experiments, Plexon Offline Sorter (Plexon, Dallas, TX), was used to remove artifacts. Principal component analysis (PCA) and waveform shape were used for spike sorting. Single units were defined as those 1) having a consistent waveform shape, 2) being a separable cluster in PCA space, and 3) having a consistent refractory period of at least 2 milliseconds in interspike interval histograms. The same MSNs were sorted across saline, D2 blockade, and D1 blockade sessions by loading all sessions simultaneously in Offline Sorter and sorted using the preceding criteria. MSNs had to have consistent firing in all sessions to be included. Sorting integrity across sessions was quantified by comparing waveform similarity via correlation coefficients between sessions.

Spike activity was analyzed for all cells that fired between 0.5 Hz and 20 Hz over the entire behavioral session. Putative MSNs were further separated from striatal fast-spiking interneurons (FSIs) based on hierarchical clustering of the waveform peak-to-trough ratio and the half-peak width (*fitgmdist* and *cluster.m*; Fig S1; (Berke, 2011)). We calculated kernel density estimates of firing rates across the interval (-4 seconds before trial start to 22 seconds after trial start) binned at 0.2 seconds, with a bandwidth of 1. We used PCA to identify data-driven patterns of z-scored neuronal activity, as in our past work (Bruce et al., 2021; Emmons et al., 2017; Kim et al., 2017a). The variance of PC1 was empirically compared against data generated from 1000 iterations of data from random timestamps with identical bins and kernel density estimates.

Average plots were shown with Gaussian smoothing for plotting purposes only.

### Immunohistochemistry

Following completion of experiments, mice were transcardially perfused with ice-cold 1x phosphate-buffered saline (PBS) and 4% paraformaldehyde (PFA) after anesthesia using ketamine (100 mg/kg IP) and xylazine (10 mg/kg IP). Brains were then fixed in solutions of 4% PFA and 30% sucrose before being cryosectioned on a freezing microtome. Sections were stained for tyrosine hydroxylase with primary antibodies for >12 hours (rabbit anti-TH; Millipore MAB152; 1:1000) at 4° C. Sections were subsequently visualized with Alexa Fluor fluorescent secondary antibodies (goat anti-rabbit IgG Alexa 519; Thermo Fisher Scientific; 1:1,000) matched to host primary by incubating 2 hours at room temperature. Histological reconstruction was completed using postmortem analysis of electrode placement by slide-scanning microscopy on an Olympus VS120 microscope (Olympus, Center Valley, PA; Fig S2&S4).

### Trial-by-trial GLMs

To measure time-related ramping over the first 6 seconds of the interval, we used trial-by-trial generalized linear models (GLMs) at the individual neuron level in which the response variable was firing rate and the predictor variable was time in the interval or nosepoke rate (Shimazaki and Shinomoto, 2007). For each neuron, its time-related “ramping” slope was derived from the GLM fit of firing rate vs time in the interval, for all trials per neuron. GLMs accounted for nosepoke movements unless there were no nosepokes within the 0-6 second interval. All GLMs were run at a trial-by-trial level to avoid the effects of trial averaging (Latimer et al., 2015) as in our past work (Bruce et al., 2021; Emmons et al., 2017; Kim et al., 2017b). We performed additional sensitivity analyses excluding outliers outside of 95% confidence intervals and measuring the firing rate from the start of the interval to the time of the switch response on a trial-by-trial level for each neuron.

### Machine-learning analyses

To predict time, we used a naïve Bayesian classifier to evaluate neuronal ensemble decoding as in our past work (Emmons et al., 2020, 2017; Gouvea et al., 2015; Kim et al., 2017a; Mello et al., 2015). Data from neurons with more than 20 trials across all mice with the goal of evaluating how the decoding of time is affected by the epoch in the interval and by D2/D1 blockade. To preclude edge effects that might bias classifier performance, we included data from 6 seconds before trial start and 6 seconds after the end of the interval. We used leave-one-out cross-validation to predict an objective time from the firing rate within a trial. Classifier performance was quantified by computing the R^2^ of objective time vs predicted time, only for bins during the interval (0-6, 6-12, and 12-18 seconds; see Fig 7). Classifier performance was compared using time-shuffled firing rates via a Wilcoxon signed-rank test.

### Drift-diffusion models (DDM)

We constructed a four-parameter DDM (see Equations 1-3 in Results) that accounted for the ensemble threshold behavior observed in the neural data, then used it to simulate switch response times compatible with mice behavioral data. The model was implemented in MATLAB, starting from initial value *b* and with discrete computer simulation steps 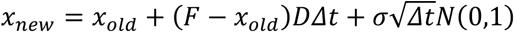. Here *N*(0,1) represents random values generated from the standardized Gaussian distribution (mean = 0 and standard deviation = 1).

The integration timestep was *Δt* = 0.1. For each numerical simulation, the “switch response time” was defined as the time *t*^∗^when variable *x* first reached the threshold value *T* = *F*(1 − *b*/4) + (1 − *F*)*b*/4. For each condition, we ran 500 simulations of the model (up to 25 seconds per trial) and recorded the switch response times. Examples of firing rate *x* dynamics are shown in Fig 4A-B. We observed that the distributions of interval timing switch response times could be fit by a gamma probability distribution function 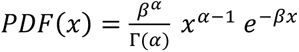 with shape *α* and rate *β* (see Fig S8 and Table S2). 10-fold cross-validation revealed highly stable fits between gamma, models and data.

### Selection of DDM parameters

Our goal was to build DDMs with dynamics that produce “response times” according to the observed distribution of mice switch times. The selection of parameter values in Fig 4 was done in three steps. First, we fit the distribution of the mice behavioral data with a Gamma distribution and found its fitting values for shape *α_M_* and rate *β_M_* (Table S2 and Fig S8; R^2^ Data vs Gamma ≥ 0.94). We recognized that the mean *μ_M_* and the coefficient of variation *CV_M_* are directly related to the shape and rate of the Gamma distribution by formulas *μ_M_* = *α_M_*/*β_M_* and 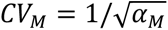. Next, we fixed parameters *F* and *b* in DDM (e.g., for D2-MSNs: *F* = 1, *b* = 0.52) and simulated the DDM for a range of values for *D* and *σ*. For each pair (*D*, *σ*), one computational “experiment” generated 500 response times with mean *μ* and coefficient of variation *CV*. We repeated the “experiment” 10 times and took the group median of *μ* and *CV* to obtain the simulation-based statistical measures *μ_S_* and *CV_S_*. Last, we plotted *E^μ^* = |(*μ_S_* − *μ_M_*)/*μ_M_*| and *E*^9:^ = |*CV_S_* − *CV_M_*|, the respective relative error and the absolute error to data (Fig S7). We considered that parameter values (*D*, *σ*) provided a good DDM fit of mice behavioral data whenever *E^μ^* ≤ 0.05 and *E*^9:^ ≤ 0.02 (Fig S7).

### Analysis and modeling of mouse MSN-ensemble recordings

Our preliminary analysis found that, for sufficiently large number of neurons (*N* > 11), each recorded ensemble of MSNs on a trial-by-trial basis could predict when mice would respond. We took the following approach: *First*, for each MSN, we convolved its trial-by-trial spike train *Spk*(*t*) with a 1-second exponential kernel *K*(*t*) = *w e*^-^*^t^*^/*w*,^ if *t* > 0 and *K*(*t*) = 0 if *t* ≤ 0 (Zhou et al., 2018; here *w* = 1 *s*). Therefore, the smoothed, convolved spiking activity of neuron *j* (*j* = 1,2, … *N*),

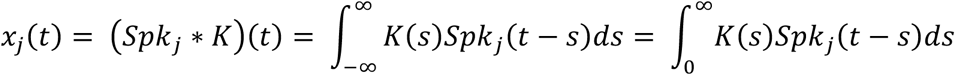

tracks and accumulates the most recent (one second, in average) firing-rate history of the *j*-th MSN, up to moment *t*. We hypothesized that the ensemble activity (*x*_1_(*t*), *x*_2_(*t*), …, *x_N_*(*t*)), weighted with some weights *β_j_*, could predict the trial switch time *t*^∗^ by considering the sum

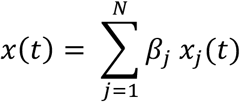

and the sigmoid

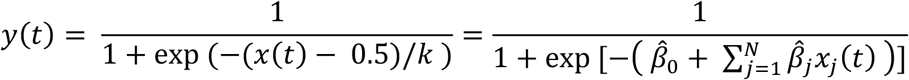

that approximates the firing rate of an output unit. Here parameter *k* indicates how fast *x*(*t*) crosses the threshold 0.5 coming from below (if *k* > 0) or coming from above (if *k* < 0) and relates the weights *β̂_j_* to the unknowns *β_j_* = *β_j_*/*k* and *β̂*_0_ = −0.5/*k*. Next, we ran a logistic fit for every trial for a given mouse over the spike count predictor matrix (*x*_1_(*t*), *x*_2_(*t*), …, *x_N_*(*t*)Z from the mouse MSN recorded ensemble, and observed value *t*^∗^, estimating the coefficients *β̂*_0_ and *β̂_j_*, and so, implicitly, the weights *β_j_*. From there, we compute the predicted switch time *t*^∗^*_pred_* by condition *x*(*t*) = 0.5. Accuracy was quantified comparing the predicted accuracy within a 1 second window to switch time on a trial-by-trial basis (Fig S4).

### Statistics

All data and statistical approaches were reviewed by the Biostatistics, Epidemiology, and Research Design Core (BERD) at the Institute for Clinical and Translational Sciences (ICTS) at the University of Iowa. All code and data are made available at http://narayanan.lab.uiowa.edu/article/datasets. We used the median to measure central tendency and the interquartile range to measure spread. We used Wilcoxon nonparametric tests to compare behavior between experimental conditions and Cohen’s *d* to calculate effect size. Analyses of putative single-unit activity and basic physiological properties were carried out using custom routines for MATLAB.

For all neuronal analyses, variability between animals was accounted for using generalized linear-mixed effects models and incorporating a random effect for each mouse into the model, which allows us to account for inherent between-mouse variability. We used *fitglme* in MATLAB and verified main effects using *lmer* in R. We accounted for variability between MSNs in pharmacological datasets in which we could match MSNs between saline, D2 blockade, and D1 blockade. P values < 0.05 were interpreted as significant.

## Data Availability

All raw data are available at http://narayanan.lab.uiowa.edu/article/datasets.

## Code Availability

All code is available at http://narayanan.lab.uiowa.edu/article/datasets.

## Acknowledgements

This work was funded by NIMH R01MH116043 to NSN. We thank editors, reviewers, Bernardo Sabatini, Filipe Rodrigues, and Joseph Paton for detailed engagement with our manuscript.

## Author Contributions

RAB, MAW, and NN designed the experiments. RAB, MAW, ASB, RV, CJ, and HRS performed all experiments and collected data. The data was analyzed by RAB, MAW, ASB, HRS, YK, RC, and NN. RC developed the computational model. RAB, MAW, ASB, KS, RC, and NN wrote the manuscript, and all authors reviewed and revised the manuscript.

## Supplementary Information

**Table S2:** Gamma distribution parameters.

| *Condition* | *Source* | $\alpha$ | $\beta$ | |
| --- | --- | --- | --- | --- |
| *D2-MSNs Laser Off* | Empirical Data | 6.03 | 0.67 | Fig S8A |
| *D2-MSNs Laser Off* | Model | 6.08 | 0.69 | Fig S8E |
| *D2-MSNs Inhibited* | Empirical Data | 7.62 | 0.78 | Fig S8B |
| *D2-MSNs Inhibited* | Model | 7.67 | 0.79 | Fig S8F |
| *D1-MSNs Laser Off* | Empirical Data | 5.87 | 0.68 | Fig S8C |
| *D1-MSNs Laser Off* | Model | 5.89 | 0.69 | Fig S8G |
| *D1-MSNs Inhibited* | Empirical Data | 7.13 | 0.70 | Fig S8D |
| *D1-MSNs Inhibited* | Model | 7.31 | 0.72 | Fig S8H |

**Figure S1:**
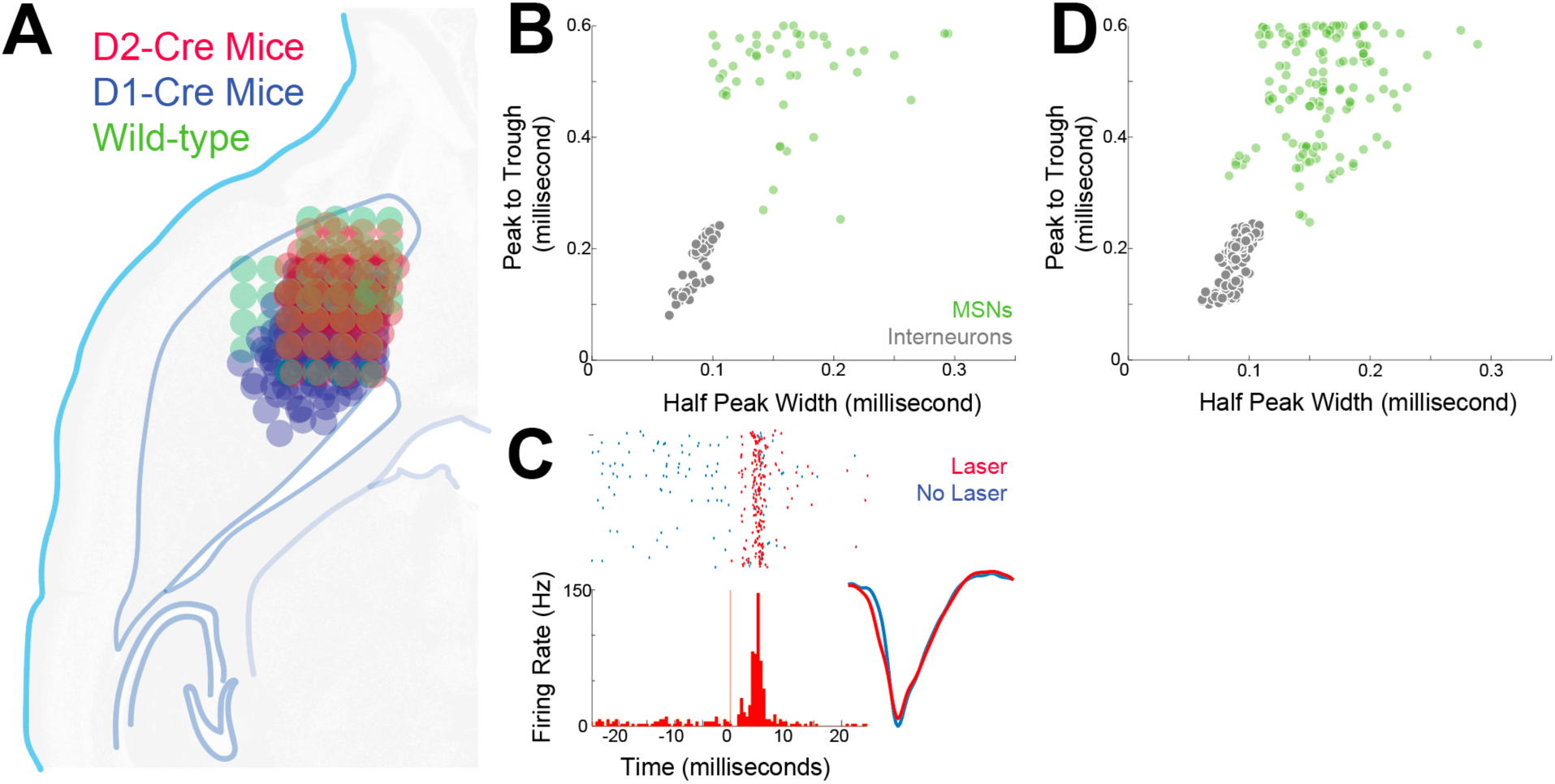
A) Recording locations in the dorsomedial striatum (targeting AP +0.4, ML -1.4, DV -2.7). Electrode reconstructions for D2-Cre (red), D1-Cre (blue), and wild-type mice (green). Only the left striatum was implanted with electrodes in all animals. B) MSN classification by waveform criteria for sessions with optogenetic tagging. C) Example of an optogenetically tagged MSN. This neuron expresses ChR2 and fired action potentials within 5 milliseconds of 473 nm laser pulses (red line). Spikes from laser trials shown as red ticks; trials without laser shown as blue ticks. Inset on bottom right – waveforms from laser trials (red) and trials without laser (blue). Across 73 tagged neurons, waveform correlation coefficients for laser trials vs trials without laser was r = 0.97 (0.92-0.99), indicating that optogenetically triggered spikes were similar to non-optogenetically triggered spikes. D) MSN classification by waveform criteria for pharmacology sessions.

**Figure S2:**
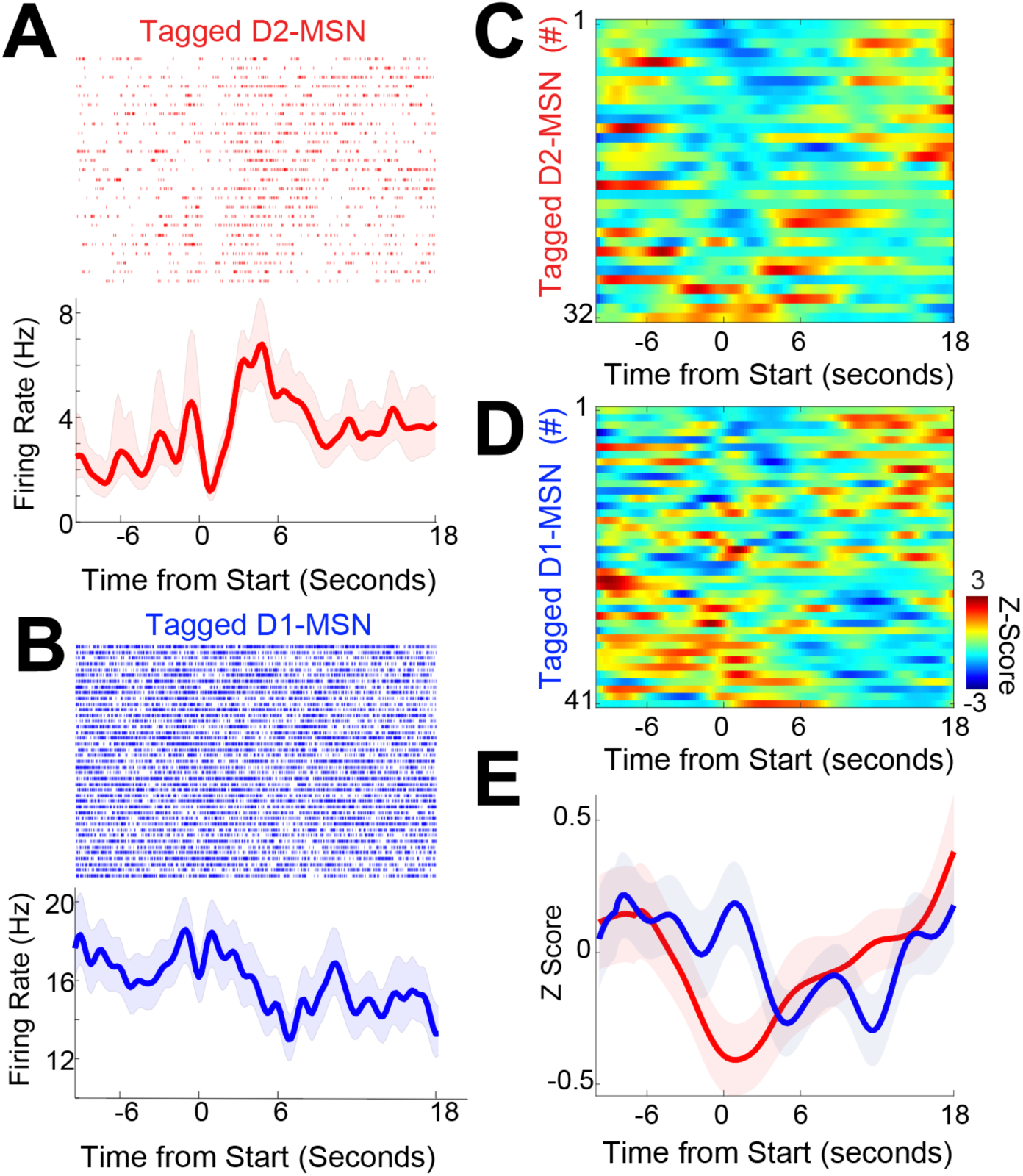
D2- and D1-MSN activity over a longer epoch from 10 seconds prior to trial start, when mice initiated trials at the back nosepoke, to the end of 18 seconds, after which making a second nosepoke led to reward. A) Tagged D2-MSN from Figure 2C shown over a longer interval, and B) tagged D1-MSN from Figure 2D shown over a longer interval. C) Peri-event time histograms from D2-MSNs and D) from D1-MSNs over a longer interval. E) We noticed that on average, D2-MSNs and D1-MSNs had the biggest differences in dynamics during the 6- second interval after trial start, where they tended to have distinct slopes (Fig 3C&D); slope analyses were less reliable for other epochs. Data from 32 tagged D2-MSNs in 4 D2-Cre mice and 41 tagged D1-MSNs in 5 D1-Cre mice as in Fig 2.

**Figure S3:**
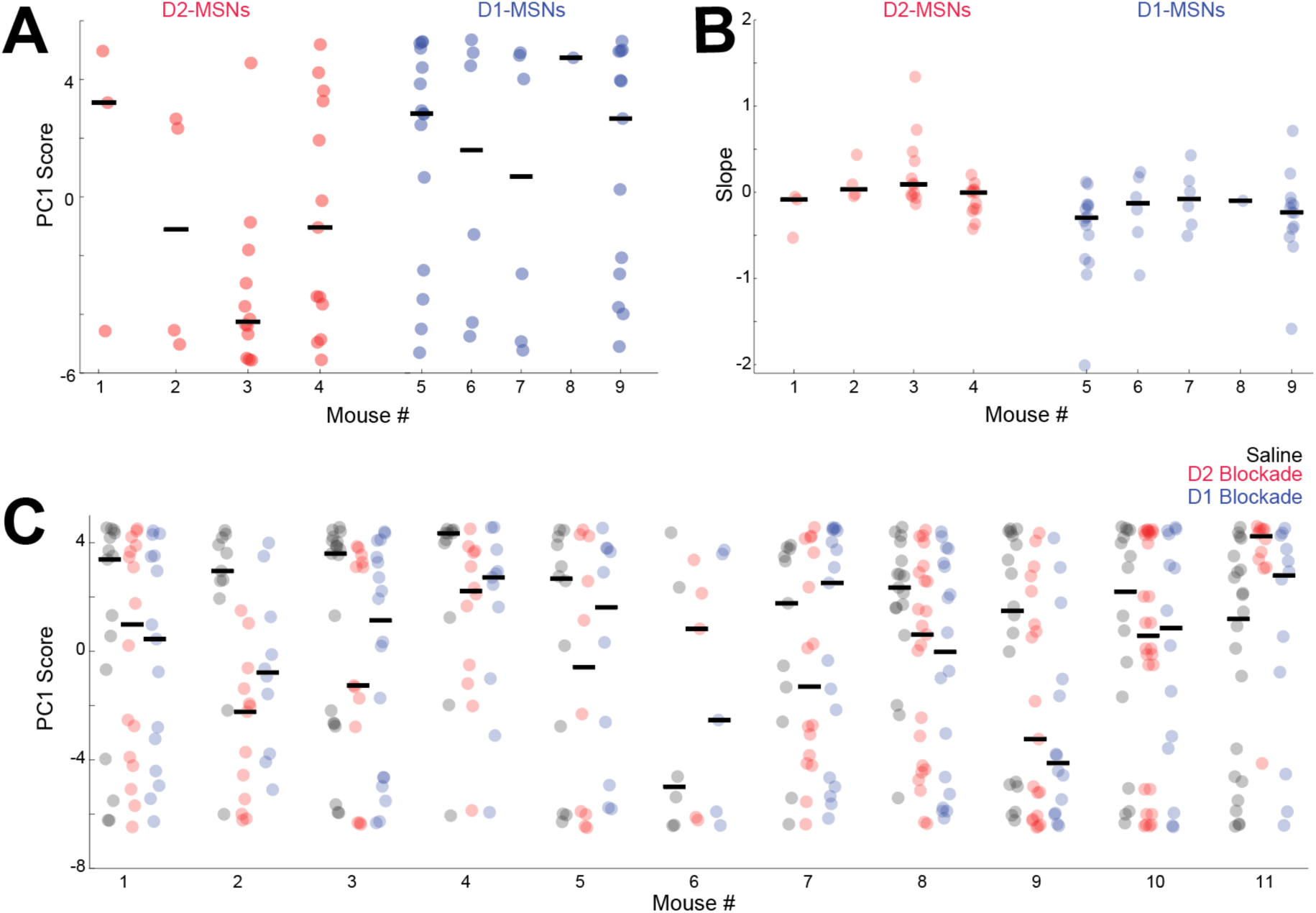
Effects in individual mice from optogenetic tagging experiments for A) PC1 and B) trial-by-trial GLM slope of firing rate over the interval; red = D2-MSN, and blue = D1-MSN. C) Effects in individual mice for PC1 for all neurons from combined pharmacology and neuronal ensemble recording; black is saline, red is D2 Blockade, and blue is D1 Blockade. Black lines indicate the median for each condition in each mouse. All statistics analyzing these data used linear-mixed effects models as incorporating a random effect for each mouse into the model allows to account for inherent between-mouse variability.

**Figure S4:**
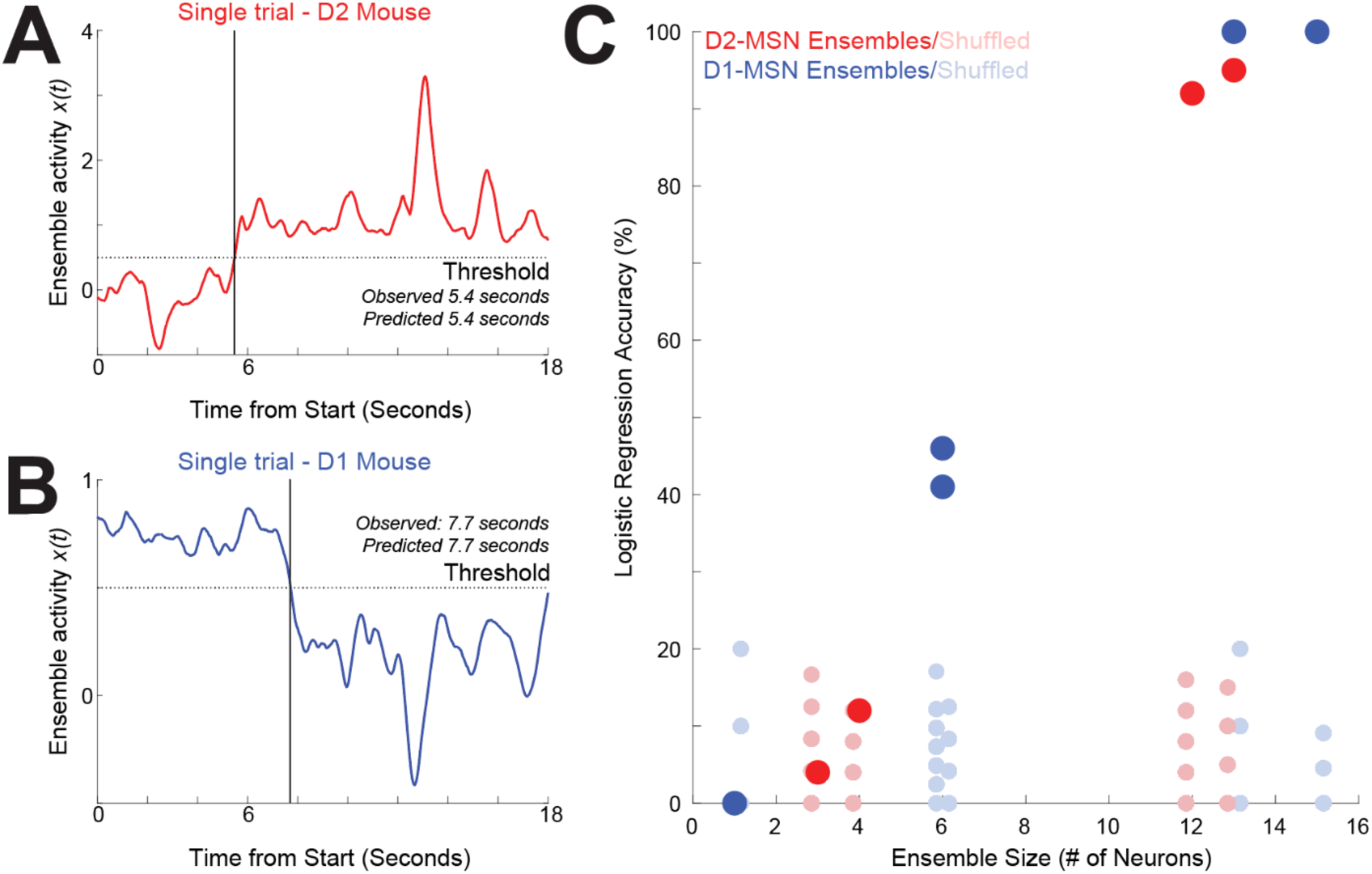
Trial-by-trial predictions of switch response time from D2-MSN and D1-MSN ensemble dynamics. A) Exemplar integral of network activity *x*(*t*) = ∑ *β_j_x_j_*(*t*) generated from an ensemble of 13 D2-MSNs for a single animal on a single trial. The coefficients *β*_!_ were computed from the logistic regression fit to the switch time *t*^∗^using the neurons firing rates (*x_j_*(*t*))*_j_* as the predictor matrix (see Methods). On this trial, this integral drifted towards the response threshold 0.5, and logistic regression accurately predicted the switch response time. B) Exemplar integral network activity *x*(*t*) generated from an ensemble of 15 D1-MSNs in a single animal on a single trial. C) Across all D2-MSN and D1-MSN ensembles per individual mouse, we found that the trial-by-trial accuracy increased with ensemble size, exceeding >90% when D2-MSN and D1-MSN ensembles were >11 neurons. Neuronal data from 5 D1- Cre (blue dots) and 4 D2-Cre (red dots) mice in Figures 2-3. Corresponding light blue/light red dots show accuracy values computed in 100 simulations of respective D2-MSN/D1-MSN ensembles with Poisson spikes matched to MSN firing rates.

**Figure S5:**
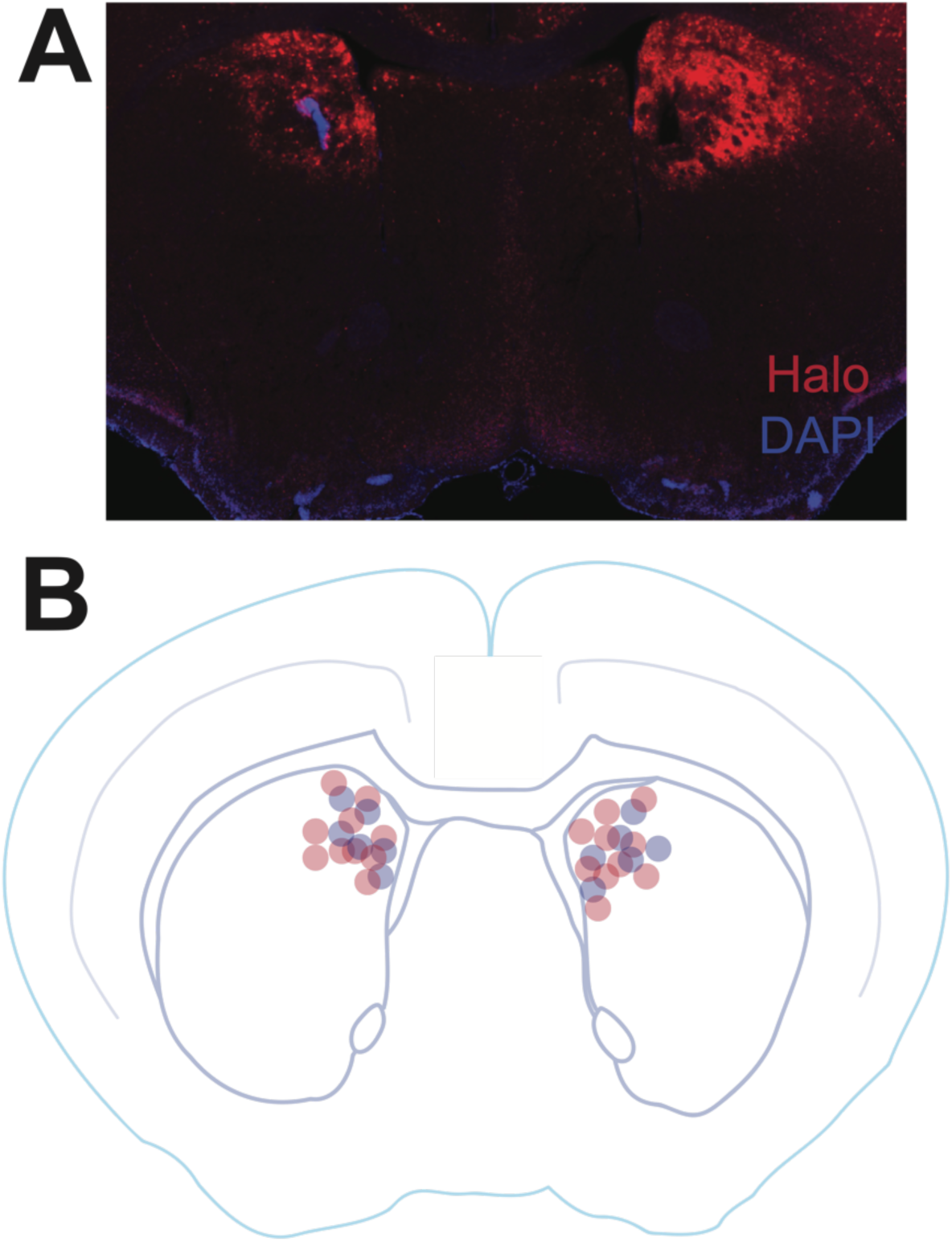
Fiber optic locations from A) an opsin-expressing mouse with mCherry-tagged halorhodopsin and bilateral fiber optics, and B) across 10 D2-Cre mice (red) and 6 D1-cre mice (blue) with fiber optics (targeting AP +0.9, ML +/-1.3, DV –2.5).

**Figure S6:**
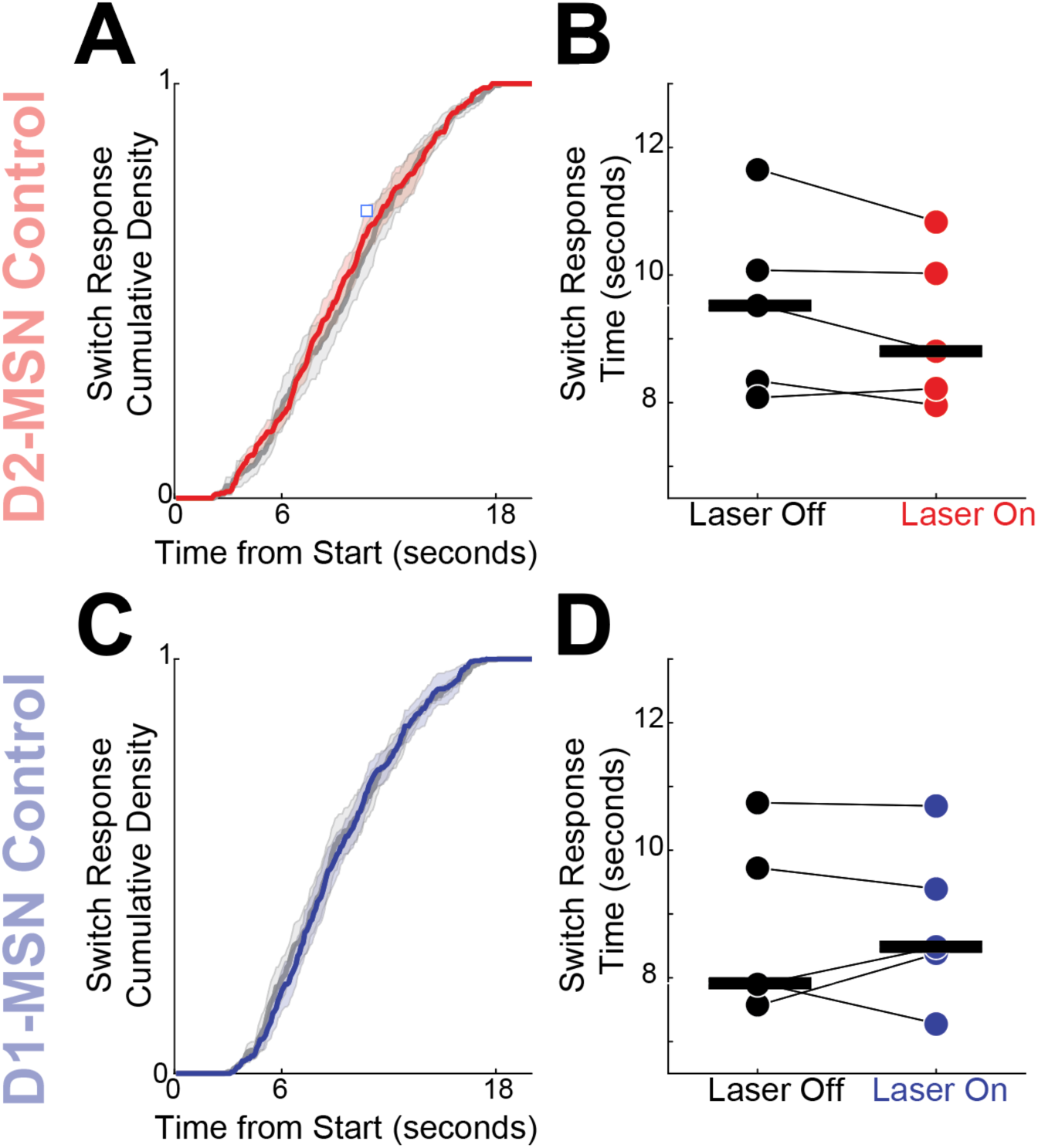
To control for heating and nonspecific effects of optogenetics, we conducted control experiments that with identical laser exposures except a virus without opsin was used. Experiments in D2-Cre mice injected with virus without opsins did not reliably affect A) cumulative density functions (CDFs) or B) switch response times (signed rank *p* = 0.44). Experiments in D1-Cre mice expressing virus without opsins did not reliably affect C) CDFs or D) switch response times (signed rank *p* = 0.81). Laser parameters (589 nm laser, 12 mW, 18 second duration) were identical to experimental animals in Fig 5.

**Fig S7:**
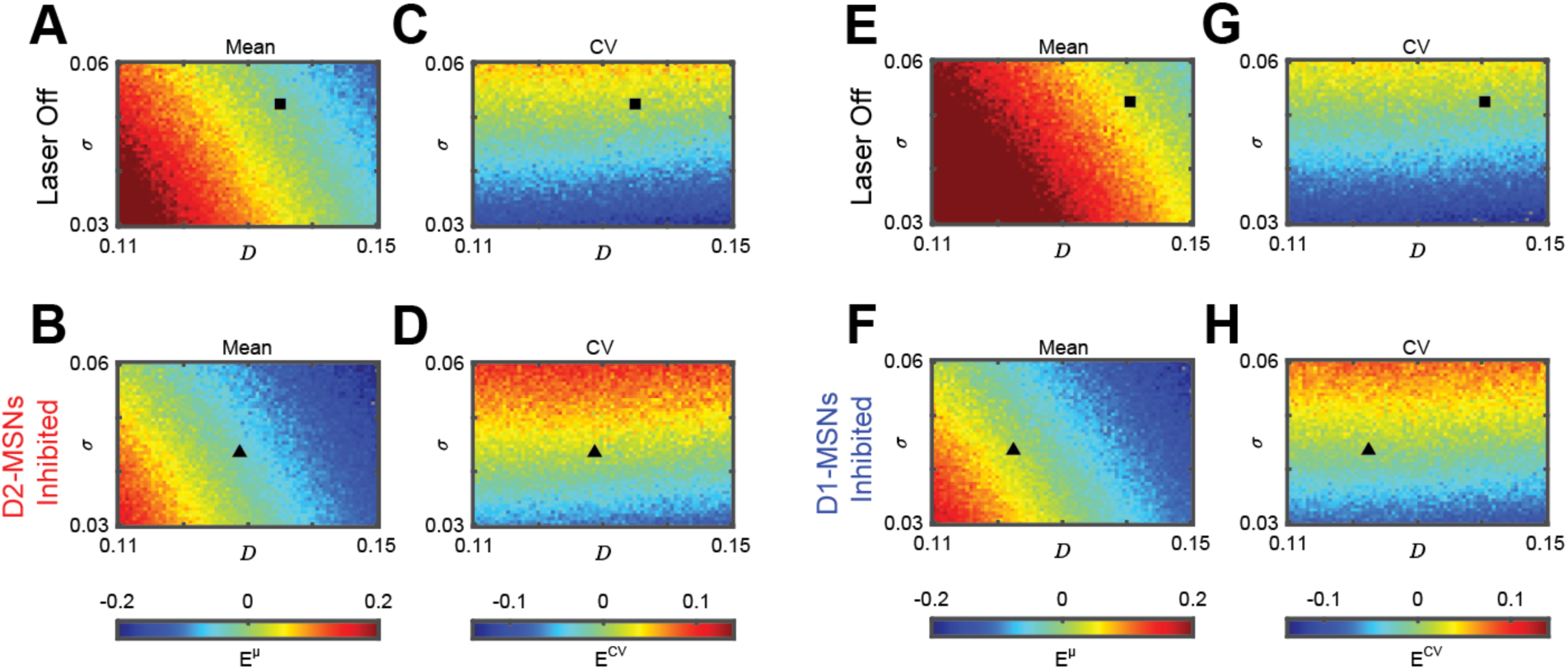
DDM parameter exploration. The relative error of simulated mean (*μ_S_*) to the behavioral gamma-fit mean (*μ_M_*), or *E^μ^* = |(*μ_S_* − *μ_M_*)/*μ_M_*|, for A) D2-Cre mice with Laser Off and for B) D2-MSN inhibition. The absolute error of the DDM computed coefficient of variation *CV_S_* relative to the behavioral Gamma-fit *CV_M_*, or (*E*^9:^ = |*CV_S_* − *CV_M_*|), for C) D2-Cre mice with Laser Off and for D) D2-MSN inhibition. The relative error of simulated mean (*μ_S_*) to the behavioral gamma-fit mean (*μ_M_*), for E) D1-Cre mice with Laser Off and for F) D1-MSN inhibition. The absolute error of the DDM computed coefficient of variation *CV_S_* relative to the behavioral Gamma-fit *CV_M_* for G) D1-Cre mice with Laser Off and for H) D1-MSN inhibition. Black squares and triangles represent parameters *D* and *σ* for Fig 4.

**Figure S8:**
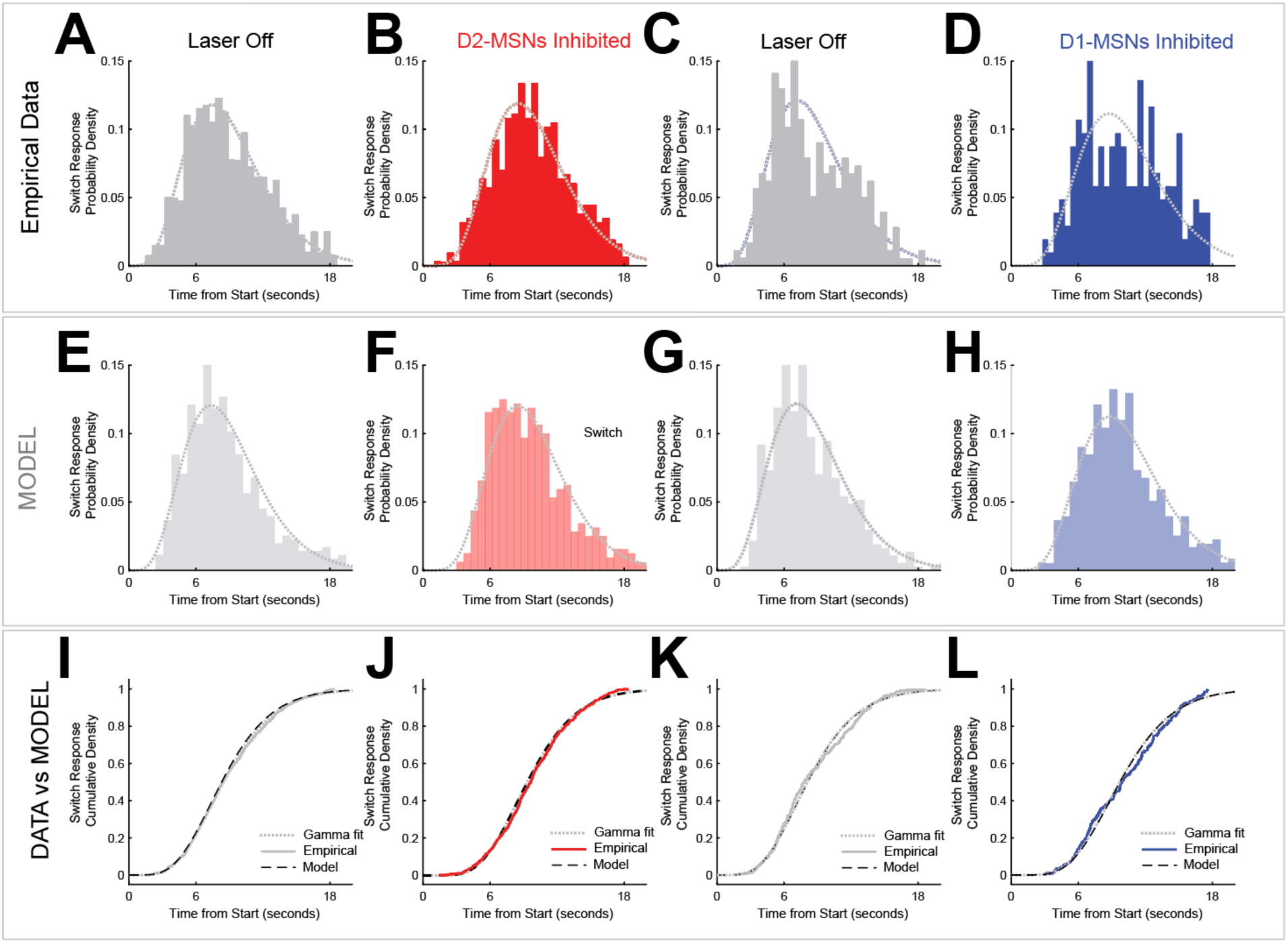
Model details. Histograms of behavioral data from D2-Cre mice with A) Laser Off and B) D2-MSN inhibition (red). Data from D1-Cre mice with C) Laser Off and with D) D1- MSN inhibition (blue). E-H) model predictions. I-L) Comparisons of empirical data vs model. All panels: fits for the gamma distribution with dotted circles; see Table S2 for the parameter values defining each gamma distribution. Behavioral data: from 10 D2-Cre mice and 6 D1-mice from Fig 5A-D. Model data: from numerical simulations of the DDM model shown in Fig 4.

**Figure S9:**
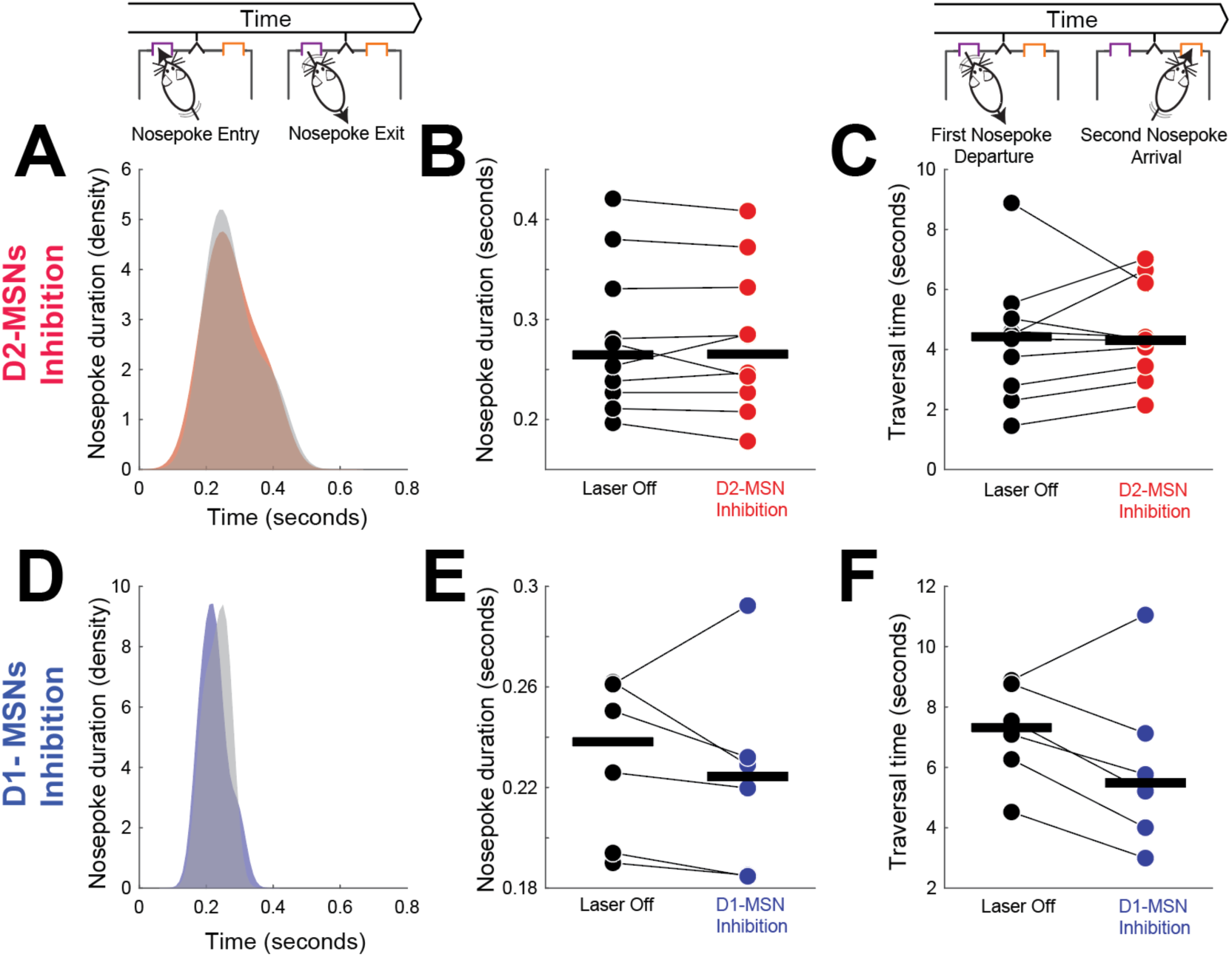
Optogenetically inhibiting D2-MSNs or D1-MSNs does not affect task-specific motor control. We measured nosepoke duration (time of nosepoke entry to exit) on switch responses. During interval timing there was no effect of optogenetic inhibition (red) of dorsomedial striatal D2-MSNs on A-B) nosepoke duration or C) the traversal time between the first and second nosepokes; traversal time is distinct from the switch response, which is the moment animals depart the first nosepoke prior to arriving at the second nosepoke. There was also no effect of optogenetic inhibition (blue) of dorsomedial striatal D1-MSNs on nosepoke duration (D-E) or F) switch traversal time. Data from the same 10 D2-Cre mice and 6 D1-Cre mice, as in Fig 5. Horizontal black lines in B, C, E, and F represent group medians.

**Figure S10:**
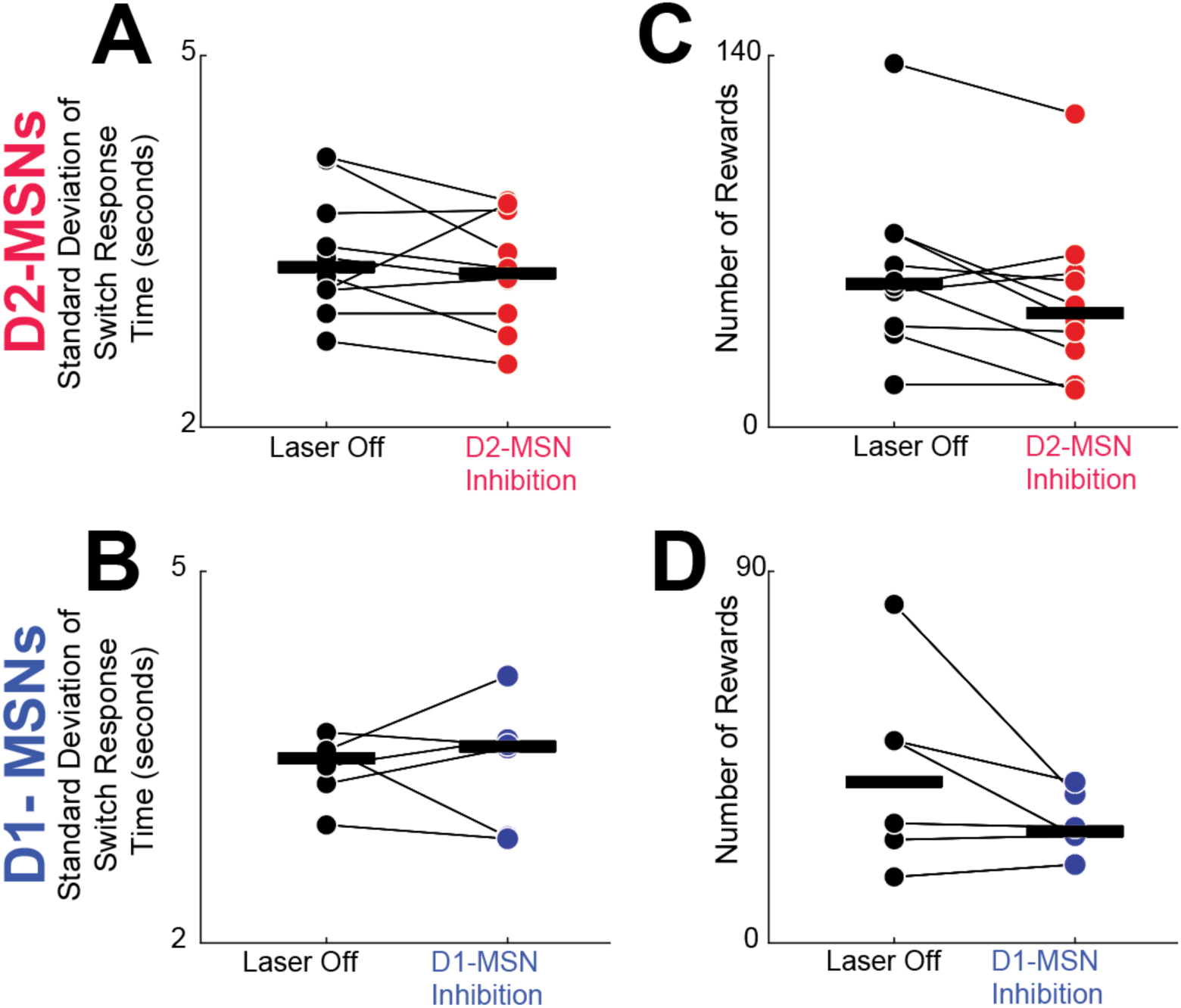
A) Standard deviation of switch response times for D2-MSN inhibition sessions (signed rank test, *p* = 0.19) and B) D1-MSN inhibition sessions (*p* = 0.84), and the number of total rewards for C) D2-MSN inhibition sessions (*p* = 0.07) and d) D1-MSN inhibition sessions (*p* = 0.25). Data from 10 D2-Cre mice and 6 D1-Cre mice as in Fig 5.

**Figure S11:**
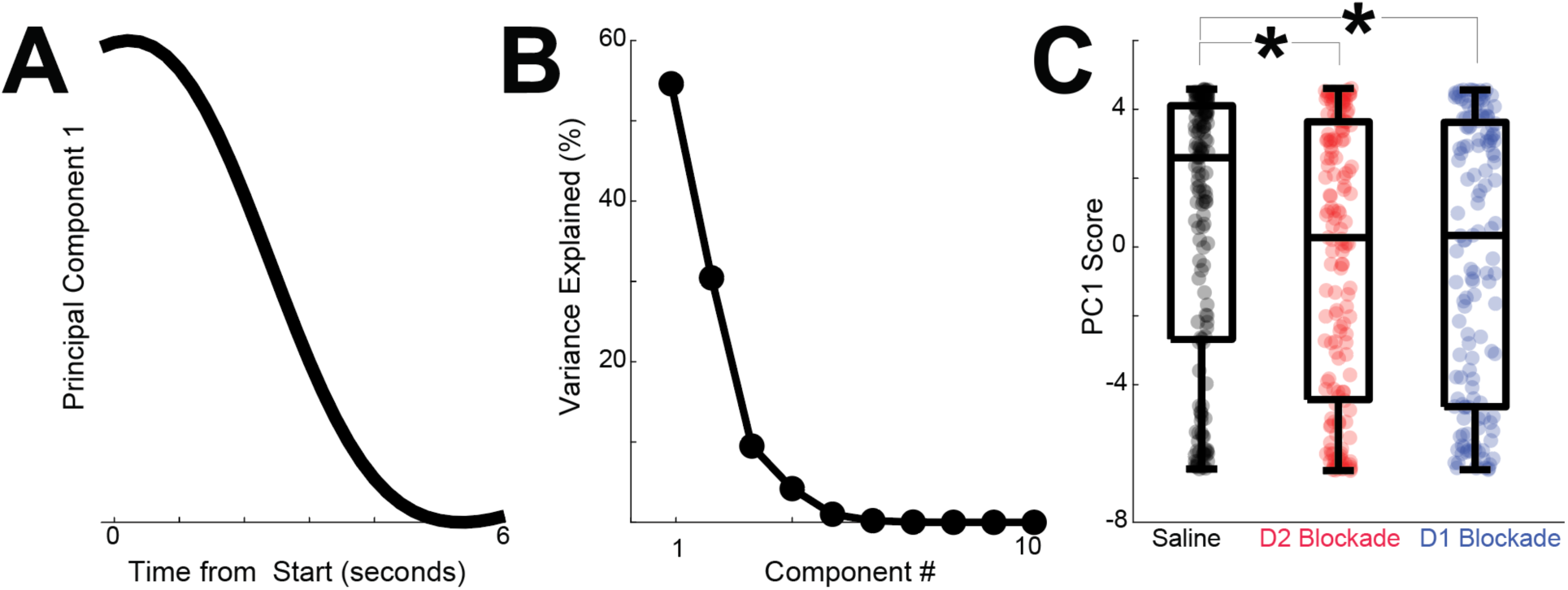
We analyzed MSN ensembles in sessions with saline (158 neurons), D2 blockade (167 neurons), or D1 blockade (144 neurons) – unlike Figure 6; all sessions were sorted independently and assumed to be fully statistically independent. A) Principal component analysis (PCA) identified MSN ensemble patterns of activity. The first principal component (PC1) exhibited time-dependent ramping. B) PC1 explained 55% of population variance among MSN ensembles. C) PC1 scores were shifted and significantly different with D2 or D1 blockade as when all sessions were sorted together in Fig 6. * = p < 0.05 via linear mixed effects models accounting for variance between mice; all analyses assumed statistical independence.

## REFERENCES

Albin RL, Young AB, Penney JB. 1989. The functional anatomy of basal ganglia disorders. Trends Neurosci 12:366–375. doi:10.1016/0166-2236(89)90074-X

Alexander GE, Crutcher MD. 1990. Functional architecture of basal ganglia circuits: neural substrates of parallel processing. Trends Neurosci 13:266–271. doi:10.1016/0166-2236(90)90107-L

Andreasen NC. 1999. Understanding the causes of schizophrenia. N Engl J Med 340:645–647. doi:10.1056/NEJM199902253400811

Averbeck BB, Lehman J, Jacobson M, Haber SN. 2014. Estimates of projection overlap and zones of convergence within frontal-striatal circuits. J Neurosci 34:9497–9505.

Balci F, Gallistel CR. 2006. Cross-domain transfer of quantitative discriminations: Is it all a matter of proportion? Psychon Bull Rev 13:636–642. doi:10.3758/BF03193974

Balci F, Papachristos EB, Gallistel CR, Brunner D, Gibson J, Shumyatsky GP. 2008. Interval timing in genetically modified mice: a simple paradigm. Genes Brain Behav 7:373–384. doi:10.1111/j.1601-183X.2007.00348.x

Berke JD. 2011. Functional properties of striatal fast-spiking interneurons. Front Syst Neurosci 5:45.

Brown SW. 2006. Timing and executive function: Bidirectional interference between concurrent temporal production and randomization tasks. Mem Cognit 34:1464–1471. doi:10.3758/BF03195911

Bruce RA, Weber MA, Volkman RA, Oya M, Emmons EB, Kim Y, Narayanan NS. 2021. Experience-related enhancements in striatal temporal encoding. Eur J Neurosci 54:5063– 5074. doi:10.1111/ejn.15344

Buhusi CV, Meck WH. 2005. What makes us tick? Functional and neural mechanisms of interval timing. Nat Rev Neurosci 6:755–765. doi:10.1038/nrn1764

Cruz BF, Guiomar G, Soares S, Motiwala A, Machens CK, Paton JJ. 2022. Action suppression reveals opponent parallel control via striatal circuits. Nature 607:521–526. doi:10.1038/s41586-022-04894-9

Cui G, Jun SB, Jin X, Pham MD, Vogel SS, Lovinger DM, Costa RM. 2013. Concurrent activation of striatal direct and indirect pathways during action initiation. Nature 494:238–242. doi:10.1038/nature11846

De Corte BJ, Wagner LM, Matell MS, Narayanan NS. 2019a. Striatal dopamine and the temporal control of behavior. Behav Brain Res 356:375–379. doi:10.1016/j.bbr.2018.08.030

De Corte BJ, Wagner LM, Matell MS, Narayanan NS. 2019b. Striatal dopamine and the temporal control of behavior. Behav Brain Res 356:375–379. doi:10.1016/j.bbr.2018.08.030

Drew MR, Fairhurst S, Malapani C, Horvitz JC, Balsam PD. 2003. Effects of dopamine antagonists on the timing of two intervals. Pharmacol Biochem Behav 75:9–15.

Drew MR, Simpson EH, Kellendonk C, Herzberg WG, Lipatova O, Fairhurst S, Kandel ER, Malapani C, Balsam PD. 2007. Transient Overexpression of Striatal D2 Receptors Impairs Operant Motivation and Interval Timing. J Neurosci 27:7731–7739. doi:10.1523/JNEUROSCI.1736-07.2007

Emmons EB, De Corte BJ, Kim Y, Parker KL, Matell MS, Narayanan NS. 2017. Rodent medial frontal control of temporal processing in the dorsomedial striatum. J Neurosci 37:8718– 8733.

Emmons EB, Kennedy M, Kim Y, Narayanan NS. 2019. Corticostriatal stimulation compensates for medial frontal inactivation during interval timing. Sci Rep 9:1–9. doi:10.1038/s41598-019-50975-7

Emmons EB, Tunes-Chiuffa G, Choi J, Bruce RA, Weber MA, Kim Y, Narayanan NS. 2020. Temporal Learning Among Prefrontal and Striatal Ensembles. Cereb Cortex Commun 1. doi:10.1093/texcom/tgaa058

Farajzadeh M, Sanayei M. 2024. Working memory affects motor, but not perceptual timing. doi:10.1101/2024.06.25.600202

Gouvea TS, Monteiro T, Motiwala A, Soares S, Machens C, Paton JJ. 2015. Striatal dynamics explain duration judgments. eLife 4. doi:10.7554/eLife.11386

Graybiel AM. 1997. The Basal Ganglia and Cognitive Pattern Generators. Schizophr Bull 23:459–469. doi:10.1093/schbul/23.3.459

Gür E, Duyan YA, Arkan S, Karson A, Balcı F. 2020. Interval timing deficits and their neurobiological correlates in aging mice. Neurobiol Aging 90:33–42. doi:10.1016/j.neurobiolaging.2020.02.021

Hinton SC, Paulsen JS, Hoffmann RG, Reynolds NC, Zimbelman JL, Rao SM. 2007. Motor timing variability increases in preclinical Huntington’s disease patients as estimated onset of motor symptoms approaches. J Int Neuropsychol Soc 13:539–543. doi:10.1017/S1355617707070671

Kim Y-C, Han S-W, Alberico SL, Ruggiero RN, De Corte B, Chen K-H, Narayanan NS. 2017a. Optogenetic Stimulation of Frontal D1 Neurons Compensates for Impaired Temporal Control of Action in Dopamine-Depleted Mice. Curr Biol CB 27:39–47. doi:10.1016/j.cub.2016.11.029

Kim Y-C, Han S-W, Alberico SL, Ruggiero RN, De Corte B, Chen K-H, Narayanan NS. 2017b. Optogenetic stimulation of frontal D1 neurons compensates for impaired temporal control of action in dopamine-depleted mice. Curr Biol 27:39–47.

Kravitz AV, Freeze BS, Parker PR, Kay K, Thwin MT, Deisseroth K, Kreitzer AC. 2010. Regulation of parkinsonian motor behaviours by optogenetic control of basal ganglia circuitry. Nature 466:622–626.

Larson T, Khandelwal V, Weber MA, Leidinger MR, Meyerholz DK, Narayanan NS, Zhang Q. 2022. Mice expressing P301S mutant human tau have deficits in interval timing. Behav Brain Res 432:113967. doi:10.1016/j.bbr.2022.113967

Latimer KW, Yates JL, Meister ML, Huk AC, Pillow JW. 2015. Single-trial spike trains in parietal cortex reveal discrete steps during decision-making. Science 349:184–187.

Matell MS, Meck WH. 2004. Cortico-striatal circuits and interval timing: coincidence detection of oscillatory processes. *Cogn Brain Res*, Neuroimaging of Interval Timing 21:139–170. doi:10.1016/j.cogbrainres.2004.06.012

Matell MS, Meck WH, Nicolelis MAL. 2003. Interval timing and the encoding of signal duration by ensembles of cortical and striatal neurons. Behav Neurosci 117:760–773. doi:10.1037/0735-7044.117.4.760

Meck WH. 2006. Neuroanatomical localization of an internal clock: A functional link between mesolimbic, nigrostriatal, and mesocortical dopaminergic systems. Brain Res 1109:93– 107. doi:10.1016/j.brainres.2006.06.031

Mello GBM, Soares S, Paton JJ. 2015. A Scalable Population Code for Time in the Striatum. Curr Biol 25:1113–1122. doi:10.1016/j.cub.2015.02.036

Merchant H, de Lafuente V. 2014. Introduction to the neurobiology of interval timing. Adv Exp Med Biol 829:1–13. doi:10.1007/978-1-4939-1782-2_1

Middleton FA, Strick PL. 2000. Basal ganglia and cerebellar loops: motor and cognitive circuits. Brain Res Rev 31:236–250. doi:10.1016/S0165-0173(99)00040-5

Monteiro T, Rodrigues FS, Pexirra M, Cruz BF, Gonçalves AI, Rueda-Orozco PE, Paton JJ. 2023. Using temperature to analyze the neural basis of a time-based decision. Nat Neurosci. doi:10.1038/s41593-023-01378-5

Narayanan NS. 2016. Ramping activity is a cortical mechanism of temporal control of action. Curr Opin Behav Sci 8:226–230. doi:10.1016/j.cobeha.2016.02.017

Narayanan NS, Albin RL. 2022. Cognition in Parkinson’s Disease, Progress in Brain Research. Elsevier, Acad. Press.

Narayanan NS, Cavanagh JF, Frank MJ, Laubach M. 2013. Common medial frontal mechanisms of adaptive control in humans and rodents. Nat Neurosci 16:1888–1897. doi:10.1038/nn.3549

Narayanan NS, Laubach M. 2009. Delay activity in rodent frontal cortex during a simple reaction time task. J Neurophysiol 101:2859–2871. doi:10.1152/jn.90615.2008

Nguyen Q-A, Rinzel J, Curtu R. 2020. Buildup and bistability in auditory streaming as an evidence accumulation process with saturation. PLOS Comput Biol 16:e1008152. doi:10.1371/journal.pcbi.1008152

Nombela C, Wolpe N, Barker RA, Rowe JB. 2016. Time on timing: Dissociating premature responding from interval sensitivity in Parkinson’s disease. Mov Disord Off J Mov Disord Soc 31:1163–1172. doi:10.1002/mds.26631

Parker KL, Chen K-H, Kingyon JR, Cavanagh JF, Narayanan NS. 2015. Medial frontal ∼4-Hz activity in humans and rodents is attenuated in PD patients and in rodents with cortical dopamine depletion. J Neurophysiol 114:1310–1320. doi:10.1152/jn.00412.2015

Parker KL, Chen K-H, Kingyon JR, Cavanagh JF, Narayanan NS. 2014. D1-Dependent 4 Hz Oscillations and Ramping Activity in Rodent Medial Frontal Cortex during Interval Timing. J Neurosci 34:16774–16783. doi:10.1523/JNEUROSCI.2772-14.2014

Parker KL, Kim YC, Kelley RM, Nessler AJ, Chen K-H, Muller-Ewald VA, Andreasen NC, Narayanan NS. 2017. Delta-frequency stimulation of cerebellar projections can compensate for schizophrenia-related medial frontal dysfunction. Mol Psychiatry 22:647–655. doi:10.1038/mp.2017.50

Parker KL, Lamichhane D, Caetano MS, Narayanan NS. 2013. Executive dysfunction in Parkinson’s disease and timing deficits. Front Integr Neurosci 7:75. doi:10.3389/fnint.2013.00075

Paton JJ, Buonomano DV. 2018. The Neural Basis of Timing: Distributed Mechanisms for Diverse Functions. Neuron 98:687–705. doi:10.1016/j.neuron.2018.03.045

Robbe D. 2023. Lost in time: Relocating the perception of duration outside the brain. Neurosci Biobehav Rev 153:105312. doi:10.1016/j.neubiorev.2023.105312

Ryan MB, Bair-Marshall C, Nelson AB. 2018. Aberrant striatal activity in parkinsonism and levodopa-induced dyskinesia. Cell Rep 23:3438–3446.

Shepherd GMG. 2013. Corticostriatal connectivity and its role in disease. Nat Rev Neurosci 14:278–291. doi:10.1038/nrn3469

Shimazaki H, Shinomoto S. 2007. A Method for Selecting the Bin Size of a Time Histogram. Neural Comput 19:1503–1527. doi:10.1162/neco.2007.19.6.1503

Shin JH, Kim D, Jung MW. 2018. Differential coding of reward and movement information in the dorsomedial striatal direct and indirect pathways. Nat Commun 9:1–14.

Simen P, Balci F, de Souza L, Cohen JD, Holmes P. 2011. A model of interval timing by neural integration. J Neurosci Off J Soc Neurosci 31:9238–9253. doi:10.1523/JNEUROSCI.3121-10.2011

Singh A, Cole RC, Espinoza AI, Evans A, Cao S, Cavanagh JF, Narayanan NS. 2021. Timing variability and midfrontal ∼4 Hz rhythms correlate with cognition in Parkinson’s disease. NPJ Park Dis 7:14. doi:10.1038/s41531-021-00158-x

Soares S, Atallah BV, Paton JJ. 2016. Midbrain dopamine neurons control judgment of time. Science 354:1273–1277. doi:10.1126/science.aah5234

Stutt HR, Weber MA, Cole RC, Bova AS, Ding X, McMurrin MS, Narayanan NS. 2024. Sex similarities and dopaminergic differences in interval timing. Behav Neurosci 138:85–93. doi:10.1037/bne0000577

Tecuapetla F, Jin X, Lima SQ, Costa RM. 2016. Complementary Contributions of Striatal Projection Pathways to Action Initiation and Execution. Cell 166:703–715. doi:10.1016/j.cell.2016.06.032

Toda K, Lusk NA, Watson GDR, Kim N, Lu D, Li HE, Meck WH, Yin HH. 2017. Nigrotectal Stimulation Stops Interval Timing in Mice. Curr Biol 27:3763–3770.e3. doi:10.1016/j.cub.2017.11.003

Tosun T, Gur E, Balci F. 2016. Mice plan decision strategies based on previously learned time intervals, locations, and probabilities. Proc Natl Acad Sci U S A 113:787–792. doi:10.1073/pnas.1518316113

Wang J, Narain D, Hosseini EA, Jazayeri M. 2018. Flexible timing by temporal scaling of cortical responses. Nat Neurosci 21:102. doi:10.1038/s41593-017-0028-6

Ward RD, Kellendonk C, Kandel ER, Balsam PD. 2011. Timing as a window on cognition in schizophrenia. Neuropharmacology. doi:10.1016/j.neuropharm.2011.04.014

Weber MA, Kerr G, Thangavel R, Conlon MM, Gumusoglu SB, Gupta K, Abdelmotilib HA, Halhouli O, Zhang Q, Geerling JC, Narayanan NS, Aldridge GM. 2024. Alpha-Synuclein Pre-Formed Fibrils Injected into Prefrontal Cortex Primarily Spread to Cortical and Subcortical Structures. J Park Dis 14:81–94. doi:10.3233/JPD-230129

Weber MA, Sivakumar K, Tabakovic EE, Oya M, Aldridge GM, Zhang Q, Simmering JE, Narayanan NS. 2023. Glycolysis-enhancing α1-adrenergic antagonists modify cognitive symptoms related to Parkinson’s disease. Npj Park Dis 9:1–7. doi:10.1038/s41531-023-00477-1

Yun S, Yang B, Anair JD, Martin MM, Fleps SW, Pamukcu A, Yeh N-H, Contractor A, Kennedy A, Parker JG. 2023. Antipsychotic drug efficacy correlates with the modulation of D1 rather than D2 receptor-expressing striatal projection neurons. Nat Neurosci 1–12. doi:10.1038/s41593-023-01390-9

Zhang Q, Abdelmotilib H, Larson T, Keomanivong C, Conlon M, Aldridge GM, Narayanan NS. 2021. Cortical alpha-synuclein preformed fibrils do not affect interval timing in mice. Neurosci Lett 765:136273. doi:10.1016/j.neulet.2021.136273

